# Diel metabolic plasticity of CAM photosynthesis in MD-2 pineapple (*Ananas comosus*) under contrasting tropical environments: biochemical patterns and agronomic implications

**DOI:** 10.64898/2026.05.27.728258

**Authors:** J. Vásquez-Jiménez, D.P Bartholomew, H. Trimino-Vásquez, L.R. Villegas-Peñaranda, B. Vargas-Leitón, G. Esquivel-Hernández

## Abstract

The current understanding of Crassulacean Acid Metabolism (CAM), including semi-controlled studies in pineapple, does not fully explain outcomes observed under commercial field conditions. Although empirical agronomy confirms a strong climatic influence on growth and development, mechanistic explanations at the metabolic level—particularly for photosynthate allocation—remain scarce. This study evaluated how environmental variation affects diel CAM outputs and how such effects can be agronomically interpreted. MD-2 pineapple plants were cultivated in contrasting natural environments across Costa Rica. Leaf samples were collected at defined phenological stages and at the end of CAM Phases I and IV. Field data revealed distinct metabolic balances between soluble sugar accumulation and nocturnal malic acid content. Under high radiation and temperature, sucrose concentrations increased markedly, reflecting shifts toward leaf growth over stem reserve storage. These shifts were associated with differences in harvest index, highlighting the role of sucrose dynamics in phenotypic plasticity. From a seed selection perspective, integrating CAM diel profiling into research protocols—together with physiological age (thermal units)—could provide a stronger basis for classifying planting material beyond current fresh-weight standards. Such integration would improve the prediction of photosynthetic performance, early establishment success, and ultimately, crop uniformity at harvest. Approximately 38% of diel metabolic patterns deviated from the classical CAM model, indicating dynamic regulatory mechanisms under field conditions. Understanding these patterns could improve the interpretation of yield variability, natural flowering incidence, and harvest index across agroecosystems. Recognizing these CAM particularities offers a path to bridge the gap between fundamental CAM biochemistry and agronomic application, enabling the development of precise management strategies that respond to metabolic plasticity under real-world conditions. Closing this gap is essential to enhance productivity, uniformity, and sustainability in pineapple agroecosystems.

## 1. Introduction

Pineapple (*Ananas comosus*) is the third most important tropical fruit worldwide by production volume (Leal & Coppens d’Eeckenbrugge, 2018), with Costa Rica leading global exports of the MD-2 cultivar, which accounts for over 80% of the international fresh pineapple trade (FAO, 2023; Sanewski et al., 2018).

As a constitutive Crassulacean Acid Metabolism (CAM) species, pineapple displays a characteristic diel cycle of nocturnal CO_2_ fixation and organic acid accumulation (Bartholomew, 2018). While CAM confers enhanced water-use efficiency through temporal separation of carbon assimilation, the physiological complexity and environmental responsiveness of pineapple CAM complicate its agronomic characterization.

Despite extensive biochemical and physiological research on CAM since the late 20th century (Black & Osmond, 2005), a persistent gap exists between mechanistic insights and understanding of CAM’s functional role in crop productivity under tropical field conditions. This gap is largely due to most studies focusing on non-commercial species or controlled environments that inadequately capture the metabolic plasticity and environmental sensitivity observed in pineapple. Contrary to the classical view of CAM as a fixed and uniform metabolic process, our research reveals that pineapple CAM exhibits significant diel metabolic plasticity modulated by environmental variables such as temperature and solar radiation. This plasticity influences the dynamics of key soluble carbohydrates—sucrose, glucose, fructose—and malic acid, which are closely linked to carbon allocation patterns fundamental for biomass accumulation and the transition to natural flowering. Such metabolic flexibility poses practical challenges and opportunities for crop management that remain insufficiently understood.

Our study was conducted across three experimental sites with contrasting environments: a cool, low-solar-radiation zone, and two warmer zones typical of commercial pineapple production—one representing optimal humid tropical conditions and another characterized by high temperature and solar radiation in a dry tropical setting. This design enabled a rigorous evaluation of CAM expression and metabolic responses under field-relevant conditions.

To isolate the effects of temperature and solar radiation, water availability was uniformly maintained via irrigation, ensuring water deficit did not confound results. Under these well-watered conditions, CAM intensity varied markedly among sites. The cool, low solar radiation zone displayed distinct metabolic profiles, reflecting the limiting influence of lower temperature and solar radiation on CAM function. In contrast, in the warmer zones, the relationship between CAM intensity and harvest index proved non-linear, underscoring the complexity of translating photosynthetic activity into agronomic performance.

We hypothesized that pineapple CAM metabolism exhibits distinct diel carbohydrate and organic acid profiles modulated by environmental factors, which directly influence carbon allocation strategies linked to growth and flowering. Our findings confirm metabolic plasticity that challenges these models and reveals the complex interplay between environment and physiology in crop performance.

These insights underscore the necessity of understanding how temperature and solar radiation regulate CAM-associated carbohydrate metabolism to develop agronomic strategies that mitigate abiotic stresses and optimize pineapple yield under evolving tropical conditions. Furthermore, the metabolic patterns and environmental interactions characterized here inform and shape future research directions and practical applications in pineapple cultivation.

By bridging biochemical CAM models with agronomic outcomes through metabolite profiling in realistic tropical environments, this study advances foundational knowledge essential for more informed crop management decisions and sustainable pineapple production. While our findings narrow the gap between the traditionally “closed” biochemical understanding of CAM and its complex agronomic expression, further research remains necessary to fully integrate metabolic plasticity within field-based productivity frameworks.

## 2. Materials and methods

### Experimental sites and planting design

Three experimental sites in Costa Rica were selected to represent contrasting temperature and solar radiation conditions. One of these sites corresponded to the typical environment for high pineapple productivity in the country. Altitudinal variation was used to establish the desired climatic contrasts, with sites located at 1573, 90, and 174 meters above sea level. For practical reference, these sites were designated as the Cool Zone (CZ), Warm Zone (WZ), and Typic Zone (TZ), respectively.

To introduce intra-site climatic variability, three planting dates were implemented at each location: January (S1), May (S2), and September (S3) of 2021. While the study zones differed ecologically, both WZ and TZ were of agronomic relevance. The TZ site reflects the standard Costa Rican condition for high pineapple yields. In contrast, although commercial cultivation has declined in WZ-type environments in Costa Rica, such conditions remain common in other pineapple-producing countries such as Mexico and Colombia, among others. Furthermore, climate change may shift the typical TZ environment toward WZ-like conditions in the near future.

Pineapple was not cultivated commercially in the CZ site in Costa Rica. Its inclusion in the study served an experimental purpose: to assess how low temperatures affected both the quantity and composition of photosynthates produced, as well as biometric growth characteristics.

All plants were grown in culture bags using a uniform soil substrate, aiming to minimize variability unrelated to the primary environmental factors under study (temperature and solar radiation). Across all experimental plots, the same nutritional and phytosanitary protocol was applied, following Costa Rica’s good agricultural practices for pineapple. Irrigation was standardized at 43.1 mm per week.

Each experimental plot consisted of 40 plants. For each of the three planting dates (S1, S2, S3) within each zone, one plot was established—resulting in 120 plants per zone and a total of nine plots across all sites (3 zones × 3 plantings), encompassing 360 experimental plants. This design ensured controlled comparisons across both spatial and temporal environmental gradients.

### Collecting samples in the field

Sampling was conducted when MD-2 pineapple plants reached specific vegetative (V-Stages) and reproductive (R-Stages) phenological phases, following the protocol described by Vásquez-Jiménez et al. (2023). Samples for sugar and organic acid analyses were collected at the following stages, each with three biological replicates:

V-Stages – First New Leaf (FNL), Leaf Cycle 1 (LC1), Leaf Cycle 2 (LC2), and Leaf Cycle 3 (LC3); R-Stages – Open Heart (OH), Final Anthesis (FA), and Harvest (HRV).

Once a target phenological stage was reached, three homogeneous plants were randomly selected under uniform management conditions. In each plant, the youngest C leaf was identified— typically the 13th leaf when counted backward from the smallest visible leaf in the apical whorl (Vásquez-Jiménez et al., 2023). To ensure consistent environmental exposure, the sampled leaves were oriented in the same direction across all replicates, minimizing variability due to light interception and convective cooling.

A 10 cm section was marked immediately distal to the sub-chlorophyllous region, extending toward the apex. This region represents fully developed chlorophyllous tissue, located well above the leaf base, and corresponds to the widest portion of the leaf blade, which contributes significantly to whole-plant photosynthate production (Sideris & Krauss, 1936; Vásquez-Jiménez & Bartholomew, 2018). Samples were collected at two time points aligned with CAM phases:

- H1 (sunset of Day 0): end of Phase IV of CAM, when high sugar and low malic acid levels are expected.
- H2 (dawn of Day 1): end of Phase I of CAM, when malic acid content peaks and sugar levels are low.

Both samplings were performed under low-light conditions (solar radiation < 50 W·m^−2^).

For H1, the marked 10 cm segment was excised and a longitudinal cut was made along the midline of the leaf blade. The tissue was then halved horizontally to obtain one half of the marked section. For H2, a cross-section was taken from the remaining tissue left on the plant. All samples were immediately immersed in liquid nitrogen and stored until laboratory processing.

Each time point (H1 and H2) consisted of three representative and homogeneous individuals per phenological stage, per experimental zone, and per sowing date.

The sampling process required approximately 13 months for each planting, due to the need to cover all targeted phenological stages across the full length of the pineapple crop cycle—especially in the WZ and TZ, where development is prolonged. In the CZ, slower plant growth permitted harvest-stage sampling only in the S1 planting.

### Receipt and handling of samples in the laboratory

Upon arrival at the laboratory, samples were placed in 50 mL Falcon tubes and weighed to determine fresh mass. They were then freeze-dried for a minimum of 48 hours, after which dry mass was recorded. Given the potential variation in water content and tissue density among sites, all biochemical results were expressed on a dry-weight basis. Freeze-dried tissues were stored at – 80°C until analysis. For metabolite extraction, the samples were finely ground using a mechanical mill.

### Extraction of metabolites

Metabolite extraction (sugars and organic acids) was conducted using a modified version of the sugar extraction method described by Carnal & Black (1989). Precisely 200 mg of finely ground, freeze-dried tissue was weighed to the nearest 0.1 mg and transferred to a 15 mL screw-cap tube containing 4 mL of 80% ethanol. The sample was vortexed thoroughly and then homogenized for 2 minutes at 11,000 rpm using an ULTRA-TURRAX T18 digital homogenizer. The resulting mixture was placed in a BRANSON 8510 ultrasonic bath at 55°C for 5 minutes. After rapid cooling, the tube was centrifuged at 3500 × g for 10 minutes at 4°C using a Cence H2050R refrigerated centrifuge. The supernatant was carefully transferred to a labeled 50 mL Falcon tube. The remaining pellet was subjected to three additional extractions under the same conditions. All resulting supernatants were pooled and adjusted to 25 mL with 80% ethanol in a volumetric flask. The adjusted extract was returned to the 50 mL tube and stored at −80°C until HPLC analysis.

### Total glucans analysis

Total glucans were determined using the anthrone method described by Hassid & Neufeld (1964), as applied to the ethanol-insoluble residue obtained after four successive extractions. This fraction excludes soluble sugars and includes starch plus low molecular weight glucans, as originally defined by Carnal & Black (1989). Not all samples were analyzed for this parameter. Results are reported as micro equivalents of hexose per gram of dry weight (μeq·g^−1^ DW).

### HPLC analysis

Filtered sugar extracts were transferred to 2 mL vials using 25 mm, 0.22 μm membrane filters. Analyses were conducted on a Thermo Ultimate 3000 HPLC system coupled with a Charged Aerosol Detector (CAD). For sugar quantification, a 150 mm × 3 mm amino column (3 μm particle size) was used at a flow rate of 1.0 mL·min^−1^. The column oven was maintained at 35°C, and the CAD evaporator temperature at 50°C. The mobile phase consisted of acetonitrile (eluent A) and 0.1% formic acid in water (eluent B), used in a binary gradient: 85% A from 0 to 1 min, decreasing to 50% A at 11 min, held constant until 12 min, then ramped back to 85% A at 15 min.

To compare sugars as precursors of phosphoenolpyruvate (PEP) and malic acid, sucrose, glucose, and fructose concentrations were expressed as micro equivalents of hexose per gram of dry weight (μeq·g^−1^ DW). The limits of detection (LoD) and quantification (LoQ) were as follows: sucrose, LoD = 10.7 μeq, LoQ = 35.2 μeq; glucose, LoD = 6.6 μeq, LoQ = 21.8 μeq; fructose, LoD = 7.5 μeq, LoQ = 24.7 μeq.

Organic acid (malic acid) analysis was performed using the same HPLC system equipped with a 250 mm × 4.6 mm C18 column (5 μm particle size) under isocratic elution with 0.1% formic acid in water. The flow rate was 0.7 mL·min^−1^, and both the column oven and evaporator were maintained at 35°C and 50°C, respectively. Malic acid concentrations were expressed in micromoles per gram of dry weight (μmol·g^−1^ DW). One micro equivalent of hexose is assumed to yield two micromoles of malic acid. The method’s LoD for malic acid was 41.6 μmol, and the LoQ was 137.2 μmol.

### Carbon isotopic analysis

The carbon isotope composition (δ^13^C) of 21 pineapple flesh samples was determined using a Total Organic Carbon analyzer (Aurora 1030W, OI Analytical, USA) coupled to a Cavity Ring-Down Spectrometer (CRDS, G2201-I, Picarro, USA). Approximately 50 mg of solid sample was combusted under a stream of ultra-high-purity oxygen to convert organic carbon into CO_2_. Raw δ^13^C values were obtained from the TOC-CRDS system and calibrated using internationally recognized IAEA standards (IAEA-CO-8 and IAEA-603) and in-house standards—potassium hydrogen phthalate (KHP, −26.5‰) and L-glutamic acid (−13.7‰). Results are reported in delta notation (‰) relative to the Vienna Pee Dee Belemnite (VPDB) standard.

### Statistical analysis

The experiment followed a completely randomized multi-site design. Data were analyzed using SAS software, adhering to the recommendations for analyzing agronomic experiments across multiple environments (Moore & Dixon, 2015). A fixed-effects ANOVA was employed, incorporating zone, planting season, phenological stage, sampling time, and relevant two-way interactions as fixed factors. In certain cases, the dataset was unbalanced due to the absence of LC2 and LC3 stages at the CZ site— caused by natural flowering—and the lack of harvest-stage data for some plantings in this zone, given the prolonged growth cycle.

ANOVA assumptions were verified graphically, confirming normality, independence, and homoscedasticity. Mean comparisons were conducted using the Pdiff pairwise difference test (SAS, 2019).

The statistical analysis generated 5,995 comparisons. Exploratory graphs were constructed to evaluate each metabolite by zone, planting season, phenological stage, and sampling time. Based on these visualizations, a subset of data was selected to improve agronomic interpretation of CAM metabolism in pineapple. This refined information aims to guide future research on both fundamental aspects of CAM and its agricultural application. For clarity in result interpretation—particularly regarding the balance between hexose and malic acid—selected data were synthesized into summary tables.

## 3. Results

CAM-related physiological variables—including malic acid, soluble sugars (sucrose, glucose, and fructose), and total glucans—were analyzed across all combinations of ecological zones (CZ, WZ, TZ), planting dates (S1, S2, S3), phenological stages (FNL, LC1, LC2, LC3, OH, FA, HRV), and sampling times (H1 and H2). The presentation of results follows a structure by variable, emphasizing the most informative contrasts relevant to agronomic interpretation. Particular attention is given to patterns in carbon allocation, hexose accumulation, and malic acid dynamics that illustrate the functional behavior of CAM metabolism in pineapple.

In addition to these integrated patterns, a specific analysis of individual metabolite profiles— particularly for soluble sugars (sucrose, glucose, fructose) and malic acid—is presented to identify compound-specific responses that may not be evident within the aggregate CAM configurations. Additionally, δ^13^C isotopic composition was analyzed in fruit samples from each zone as an integrative indicator of photosynthetic carbon assimilation. This approach assumes that the δ^13^C signature in the fruit reflects the carbon fixed and allocated by the plant both prior to and after floral induction (post-forcing). The carbon isotope data are presented at the end of this section, alongside the results for malic acid, to support a coherent interpretation of diel CAM expression in pineapple.

The structure of the Results section emphasizes a biochemical characterization of the diel metabolic patterns, providing a descriptive synthesis of the most informative contrasts. This framing allows the subsequent Discussion to develop a more integrative physiological and agronomic interpretation of the findings.

### Hexose content (sum of sucrose, glucose and fructose)

Hexose content ranged from just under 300 to slightly over 2150 μeq·g^−1^ dry weight. This variation is consistent with classical CAM physiology, characterized by peak hexose concentrations at dusk (H1) and minima at dawn (H2), reflecting their nocturnal utilization via glycolysis to generate phosphoenolpyruvate (PEP), as described by Carnal & Black (1989).

As this study provides the first dataset on metabolite accumulation in pineapple leaves for agronomic purposes, Figure 2 establishes a reference framework for the observed range of hexose content across all fixed effects of the experiment, including sampling times and phenological stages. Although previous studies have examined the same metabolites (Christopher & Holtum, 1996; Christopher & Holtum, 1998; Medina et al., 1993; Popp et al., 2003), their primary focus was eco-physiological rather than agronomic. Nonetheless, some of their findings offer valuable points of comparison and will be addressed later in the discussion.

**Figure 1.**
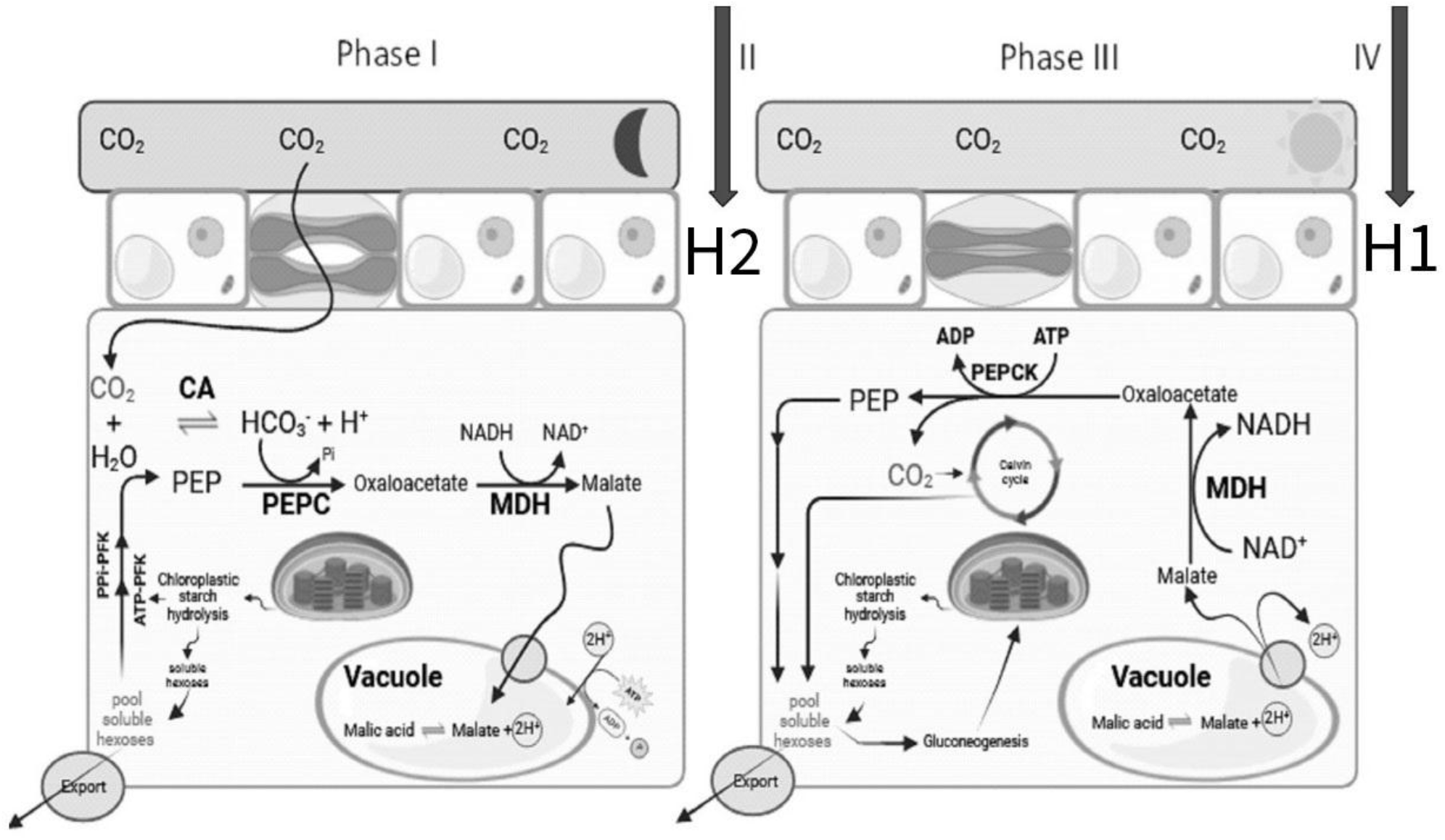
Schematic representation of CAM metabolism in pineapple throughout the 24-hour cycle. The diagram includes both nocturnal and diurnal phases and highlights the sampling points used in this study: H1 (end of Phase IV) and H2 (end of Phase I), corresponding to key transitions in malic acid, soluble sugar, and glucan dynamics. Abbreviations used in the figure: CA, carbonic anhydrase; PEP, phosphoenolpyruvic acid; PEPC, phosphoenolpyruvate carboxylase; PEPCK, phosphoenolpyruvate carboxykinase; MDH, malate dehydrogenase; PPi-PFK, pyrophosphate-dependent 6-phosphofructokinase; ATP-PFK, ATP-dependent 6-phosphofructokinase.

**Figure 2.**
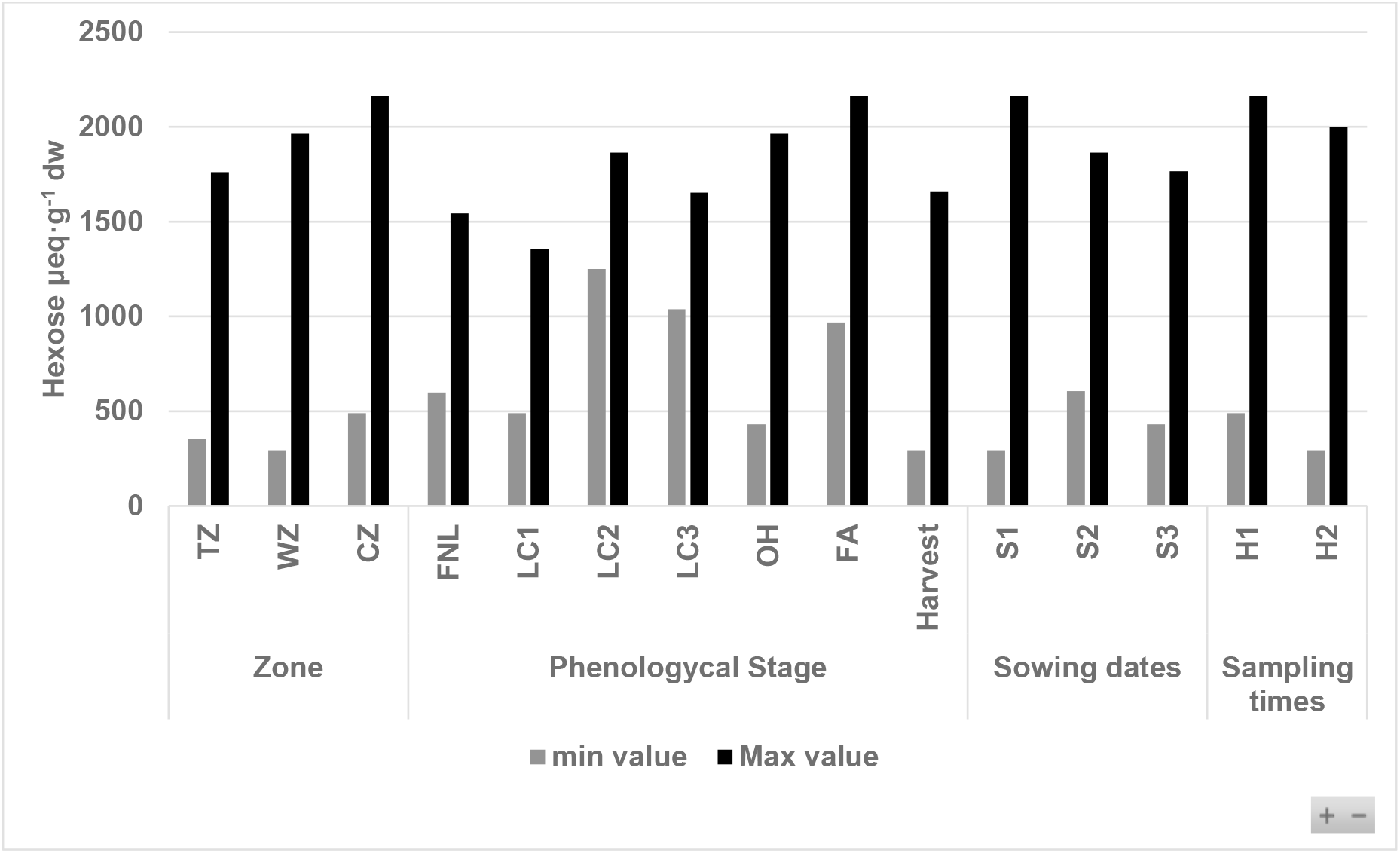
Minimum and maximum hexose contents (sum of sucrose, glucose, and fructose) measured in MD-2 pineapple leaves, categorized by fixed factors of the experimental design: ecological zone, sowing date, phenological stage, and sampling time.

No consistent relationship was found between total hexose accumulation and phenological stage. High and low hexose levels were observed across different stages, indicating that phenology alone does not determine carbohydrate production. Instead, total hexose content appears more strongly influenced by climatic variability and short-term shifts in carbon allocation. While this pattern may be expected under tropical field conditions, it contrasts with the behavior of geophytes, where carbohydrate availability during autumn and winter depends almost exclusively on starch hydrolysis (Khodorova & Boitel-Conti, 2013). As discussed later, starch degradation is also a relevant and variable source of soluble sugars in pineapple, shaped by daily environmental conditions. Malic acid synthesis is similarly influenced by climate.

Throughout the study, 5,995 individual metabolite analyses were performed, of which 749 specifically corresponded to diel reconstructions of CAM metabolism, focusing on the balance between hexoses and malic acid. This dataset enabled the identification of four functional diel metabolism patterns, selected based on their internal coherence, representativeness across contrasting environmental conditions, and relative frequency throughout the experiment. A particularly relevant finding is that all observed diel profiles could be unequivocally categorized within one of these four patterns, with no outliers or ambiguous profiles emerging. This level of functional organization suggests that these patterns represent authentic expressions of metabolic plasticity within the CAM model, rather than random variation, even in the absence of formal statistical inference between groups. These patterns are characterized by distinct diel dynamics of soluble sugars and malic acid. The first corresponds to the canonical CAM condition, characterized by complete malic acid decarboxylation by H1 and a significant decline in hexose content by H2, serving as the reference behavior. The remaining three patterns represent CAM particularities, defined as metabolite profiles deviating from this canonical condition:

1. ***A significant surplus of hexoses at H2***, suggesting either limited CAM activity or alternative metabolic regulation that prevents typical nocturnal sugar depletion.
2. ***Residual malic acid at H1 and minimal changes in hexose levels between H1 and H2***, indicating incomplete decarboxylation and restricted sugar use during the night.
3. ***A slight decrease in malic acid and an increase in hexose content at H2***, an atypical pattern possibly reflecting disruptions in CAM cycling or shifts in carbon allocation.

The analytical framework used in this section follows the approach of Carnal & Black (1989). For each sampling event, hexose content at H1 (end of the light phase) is compared with that at H2 (end of the dark phase). The reduction is interpreted primarily as the use of soluble sugars for PEP synthesis and subsequent malic acid accumulation. Thus, the balance between hexose decline and malate increase provides an estimate of CAM activity and carbon allocation efficiency. For malic acid, values at H1 are expected to match the analytical detection limit, reflecting full decarboxylation during Phase III. When values exceed the quantification threshold (LoQ), the residual is subtracted from H2, as it likely represents malate carried over from the previous cycle rather than newly fixed carbon.

### CAM-classical condition: complete malate decarboxylation by dusk and hexose depletion overnight

Table 1 illustrates the typical CAM pattern characterized by complete malic acid decarboxylation by dusk (H1) and a marked depletion of hexoses from dusk (H1) to dawn (H2). This canonical condition reflects the expected diel metabolite dynamics under fully expressed CAM. Similar trends—though not always of comparable magnitude—have been documented in nearly all studies on CAM in pineapple, including Carnal & Black (1989), Christopher & Holtum (1998), Medina et al. (1993), Popp et al. (2003).

**Table 1.** CAM-classic condition in MD-2 pineapple leaves. The data correspond to the sample from TZ-S2 at LC3, representing the condition observed in 62% of all samples in this study. This pattern is characterized by the complete decarboxylation of malic acid at H1 and a decrease in hexose content at H2.

| CAM Classic Condition |  |  |  |
| --- | --- | --- | --- |
| Fractions | Units | H1 (Mean ± SE) | H2 (Mean ± SE) |
| Malic Acid | ($\mu\text{mol g}^{-1} \text{ dw}$) | 41.6 ± 0.0 d | 433.3 ± 43.4 b |
| Total sugars | ($\mu\text{eq g}^{-1} \text{ dw}$) | 1530.2 ± 134.4 a | 1292.5 ± 96.0 abc |
| Sucrose | ($\mu\text{eq g}^{-1} \text{ dw}$) | 292.9 ± 39.4 a | 277.0 ± 51.7 a |
| Glucose | ($\mu\text{eq g}^{-1} \text{ dw}$) | 423.3 ± 34.1 a | 293.5 ± 15.7 bcd |
| Fructose | ($\mu\text{eq g}^{-1} \text{ dw}$) | 814.0 ± 63.2 ab | 722.1 ± 37.3 abc |
| Total glucans | ($\mu\text{eq g}^{-1} \text{ dw}$) | 1332.6 ± 171.3 abc | 754.4 ± 209.06 d |
| Hexose surplus | ($\mu\text{eq g}^{-1} \text{ dw}$) | 599.3 | |
|  | (%) | 277.0 |  |
| Agronomic surplus | ($\mu\text{eq g}^{-1} \text{ dw}$) | 417.9 | |
|  | (%) | 105.0 |  |

Shared letters denote statistically non-significant differences ( $P > 0.05$ ). Comparisons are valid only within each metabolite and between the two sampling times (H1 and H2). Each value represents the mean ± standard deviation of three biological replicates. The hexose surplus is calculated relative to the specific malic acid accumulation observed in the corresponding sample. The agronomic surplus is estimated using the maximum malic acid concentration recorded in the experiment as a reference for optimal CAM intensity.

The total soluble sugar content at H1 was 1530.2 μeq of hexose·g^−1^ dw, while at H2 it was 1292.5 μeq of hexose·g^−1^ dw, indicating a net decrease of 237.7 μeq of hexose·g^−1^ dw. Malic acid content at H2 reached 433.2 μmol·g^−1^ dw, with complete decarboxylation observed at H1. Climatic conditions—including solar radiation, maximum temperature, and minimum temperature—from two days prior to sampling through the day of collection are provided in Line 1 of Table 2.

**Table 2.** Solar radiation, maximum, and minimum temperature two days before (−2d), one day before (−1d), and on the day of sampling (0d) for events discussed in the text. Codes indicate zone (SZ), sowing date (Sn), and phenological stage (PS).

| Reference<br>line to the<br>text | Zone-Sowing-Phenological<br>Stage | Solar radiation<br>Mj·m<sup>-2</sup>·day<sup>-1</sup> |  |  | Tmax<br>°C |  |  | Tmin<br>°C |  |  |
| --- | --- | --- | --- | --- | --- | --- | --- | --- | --- | --- |
|  |  | -2d | -1d | 0d | -2d | -1d | 0d | -2d | -1d | 0d |
| **1** | TZ-S2-LC3 | 19.8 | 11.1 | 19.3 | 30.2 | 30.1 | 30.5 | 22.8 | 22.0 | 23.1 |
| **2** | WZ-S2-LC3 | 15.6 | 18.9 | 19.7 | 32.3 | 32.3 | 33.4 | 23.1 | 22.3 | 20.6 |
| **3** | TZ-S2-LC1 | 12.9 | 13.7 | 11.8 | 29.3 | 31.2 | 28.5 | 22.4 | 22.6 | 21.4 |
| **4** | WZ-S2-LC1 | 13.4 | 16.1 | 11.9 | 30.8 | 30.7 | 29.4 | 23.1 | 21.9 | 23.0 |
| **5** | CZ-S3-LC1 | 7.2 | 7.8 | 5.0 | 20.9 | 19.6 | 19.3 | 15.6 | 14.8 | 16.1 |
| **6** | WZ-S3-HRV | 14.1 | 14.5 | 13.7 | 29.9 | 31.1 | 30.6 | 23.5 | 22.6 | 22.8 |
| **7** | CZ-S2-HRV | 15.0 | 12.0 | 7.4 | 22.5 | 21.5 | 18.7 | 13.7 | 14.3 | 12.9 |
| **8** | CZ-S3-LC1 | 7.2 | 7.8 | 5.0 | 20.9 | 19.6 | 19.3 | 15.6 | 14.8 | 16.1 |
| **9** | WZ-S3-LC3 | 19.1 | 21.4 | 19.6 | 33.6 | 34.3 | 33.6 | 24.2 | 22.4 | 23.7 |
| **10** | WZ-S1-FNL | 20.1 | 20.4 | 20.3 | 35.3 | 33.1 | 32.8 | 18.1 | 22.8 | 23.6 |
| **11** | WZ-S3-LC2 | 22.4 | 24.6 | 21.2 | 33.6 | 34.1 | 33.7 | 18.7 | 23.2 | 22.8 |
| **12** | CZ-S2-LC1 | 11.9 | 14.0 | 22.4 | 20.2 | 20.2 | 20.7 | 13.9 | 14.1 | 14.8 |
| **13** | CZ-S1-FA | 14.9 | 12.0 | 7.4 | 22.5 | 21.5 | 18.7 | 13.7 | 14.3 | 12.9 |
| **14** | WZ-S1-LC1 | 27.2 | 24.8 | 26.0 | 35.9 | 34.9 | 34.7 | 20.8 | 19.3 | 19.2 |
| **15** | WZ-S1-LC2 | 26.6 | 11.9 | 21.1 | 32.6 | 29.9 | 33.3 | 22.3 | 23.8 | 22.5 |

Abbreviations: SZ = site/zone; Sn = sowing number; PS = phenological stage.

According to stoichiometric estimates, 433.3 μmol of malic acid require approximately 216.6 μeq of hexose. The observed sugar decline included an additional 21.1 μeq of hexose, likely exported or diverted to starch or other glucan synthesis. However, this marginal excess may be considered negligible, effectively indicating a balanced exchange.

Although our experimental design does not allow precise determination of metabolite export or timing, the simultaneous decrease in soluble sugars and total glucans from H1 to H2 suggests that hexose depletion fully accounts for malate accumulation. Under these conditions, hexose export likely occurred, with an estimated value close to the observed glucan reduction (578.2 μeq of hexose·g^−1^ dw). This inference builds directly on the stoichiometric approach developed by Carnal & Black (1989), who, through comparable calculations, identified a surplus of soluble sugars.

### CAM-particularity 1: Hexose content increases at dawn (H2), indicating atypical sugar dynamics

In this condition, total hexose content at dawn (H2) exceeded that at dusk (H1), diverging from the typical CAM pattern described in the literature (Carnal & Black, 1989; Christopher & Holtum, 1998; Medina et al., 1993; Popp et al., 2003). This atypical behavior occurred in 9% of all samples, indicating it was not an isolated event.

The total soluble sugar content increased from 1037.2 μeq of hexose·g^−1^ dw at H1 to 1353.5 μeq·g^−1^ dw at H2, yielding a gain of 316.3 μeq·g^−1^ dw. Meanwhile, malic acid accumulation at H2 reached 354.8 μmol·g^−1^ dw, which would have required 177.4 μeq of hexose according to stoichiometric estimates. This suggests a net surplus of soluble sugars. A likely explanation is glucan hydrolysis, supported by the reduction in total glucans between H1 and H2 (Table 3).

**Table 3.**
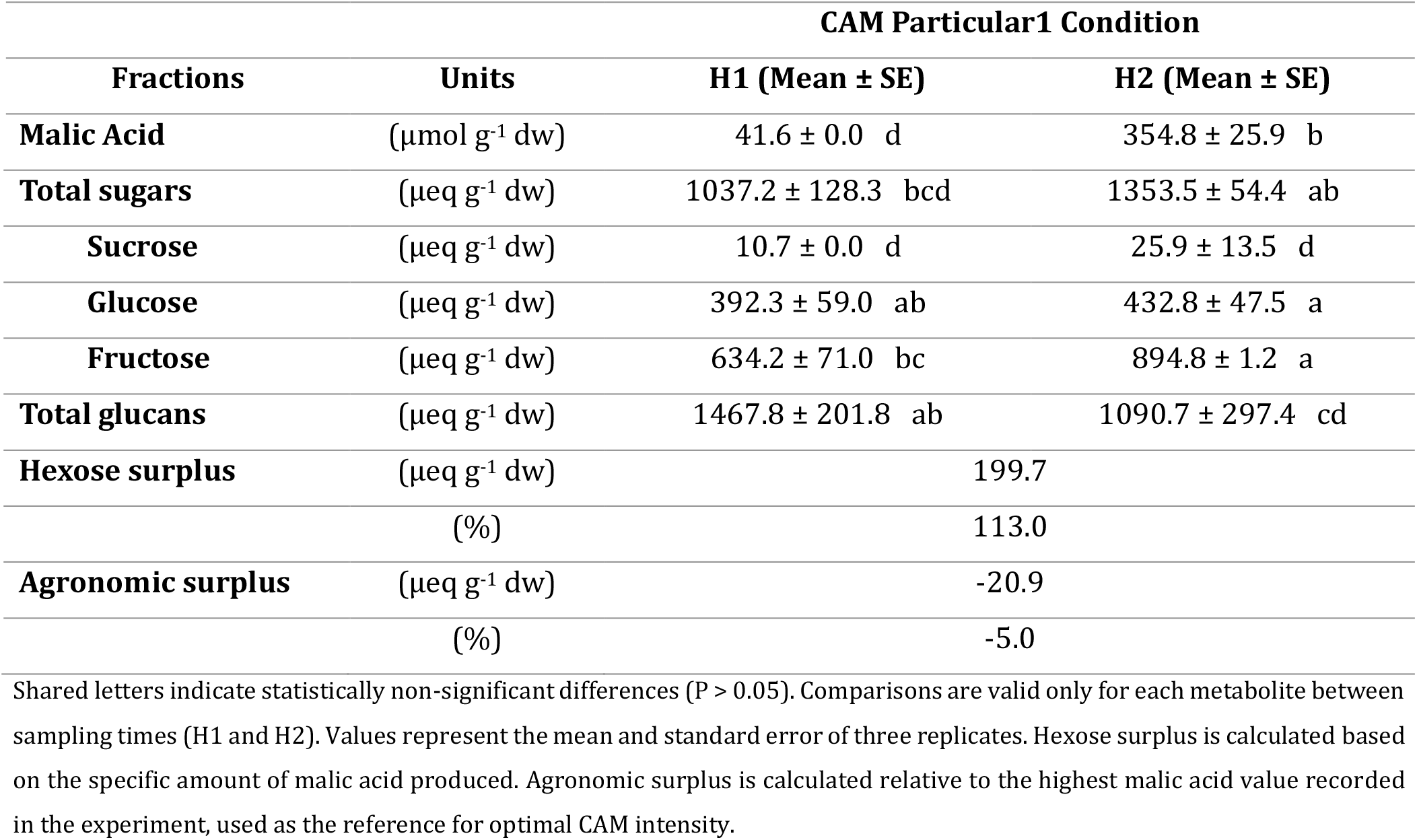
CAM-particularity 1 condition. Malic acid is completely decarboxylated at H1, while total hexose content increases at H2. (Data correspond to the sample from WZS2 in LC3 and represent the condition observed in 9% of all samples in this study).

Climatic conditions corresponding to this event are listed in Table 2, Line 2. Compared to the CAM-classical event (Line 1), the only notable difference was lower solar radiation on the day prior (-1d), which alone does not explain the change in sugar content, as similar irradiance fluctuations occurred in other events without causing this response.

These findings reinforce the idea that interpreting CAM based on 24-hour snapshots, even with frequent sampling, can be misleading. Since nocturnal carbon fixation depends on the availability of sugars derived from the previous day’s photosynthesis and starch degradation, the behavior observed in one 24-hour period is partially influenced by what occurred the day before. Thus, a full understanding of CAM requires continuous monitoring across multiple days under real field conditions, covering diverse combinations of solar radiation and temperature.

In both CAM-classical and CAM-particular1 conditions, starch hydrolysis exceeded what was required for malic acid synthesis. This implies that part of the sugars generated may have been exported. Discerning whether this exported carbon was directed toward storage or active sink tissues is critical, and techniques for distinguishing these roles—especially involving sucrose—will be discussed later.

For now, considering the findings of Carnal & Black (1989), who showed that soluble sugars serve as the substrate for PEP production in pineapple, and the general understanding that starch is stored in chloroplasts (Liao et al., 2022), both the CAM-classical and CAM-particular1 conditions imply that hexose export is linked to chloroplastic starch hydrolysis events. However, our data do not allow us to determine the timing of this export, particularly since sucrose—not hexoses—is the primary export metabolite (Hernández-Bernal et al., 2022).

In the CAM-classical condition, high amounts of hexoses at H1 were followed by a moderate decline at H2, suggesting that the export rate may have matched sucrose synthesis from starch. In CAM-particular1, sucrose levels remained low at both time points, making it harder to confirm export activity, although starch degradation still occurred. Not all starch hydrolysis led to export, as will be further explored in subsequent sections.

Although Carnal & Black (1989) reported CAM-classical behavior, our results show differences in carbon turnover. In their study, hexose turnover alone could explain nocturnal malic acid accumulation, while glucan turnover accounted for only 44%. In our CAM-classical condition, hexose turnover covered 110% of malic acid accumulation, and glucan turnover explained 167%, resulting in a combined surplus of 177%. In CAM-particular1, the increase in soluble sugars at H2 precluded calculating turnover, but the hexoses available at H1 could account for 585% of the malic acid produced during the night, and glucan turnover yielded a surplus of 113%.

These results suggest that under the conditions observed in Tables 1, 2 and 3, leaf carbohydrate reserves—including both soluble sugars and glucans—can easily support multiple rounds of nocturnal malic acid accumulation. This raises the question of whether there is a baseline threshold for hexose export, above which export occurs and below which it does not. For example, in H1 of CAM-classical, hexoses were abundant, yet starch hydrolysis still occurred. Understanding what governs this behavior could offer agronomic opportunities for managing carbon allocation in pineapple crops.

### Condition CAM-particular2: residual malic acid at H1 and minimal changes in hexoses at H2

Table 4 presents the results corresponding to CAM-particular2, observed in 19% of all samples. In this condition, malic acid was not fully decarboxylated during the day, as evidenced by the presence of significant residual malate at H1. Meanwhile, hexose levels showed minimal variation between dusk and dawn, indicating limited sugar turnover during the night.

**Table 4.** CAM-particular2 condition. Malic acid was not completely decarboxylated by H1, and hexose levels decreased only slightly by H2. (Data correspond to samples from TZS2 and WZS2 at stage LC1 and represent 19% of all samples collected in the study).

| CAM Particular 2a Condition (i.e. TZS2 in LC1) |  |  |  |
| --- | --- | --- | --- |
| Fractions | Units | H1 (Mean ± SE) | H2 (Mean ± SE) |
| Malic Acid | ($\mu\text{mol g}^{-1} \text{ dw}$) | 113.1 ± 25.2 cd | 571.4 ± 140.2 a |
| Total sugars | ($\mu\text{eq g}^{-1} \text{ dw}$) | 1301.2 ± 203.2 abc | 1223.7 ± 218.8 abc |
| Sucrose | ($\mu\text{eq g}^{-1} \text{ dw}$) | 146.2 ± 16.1 bc | 229.1 ± 56.6 ab |
| Glucose | ($\mu\text{eq g}^{-1} \text{ dw}$) | 377.2 ± 53.2 abc | 262.2 ± 60.8 d |
| Fructose | ($\mu\text{eq g}^{-1} \text{ dw}$) | 777.9 ± 135.7 ab | 732.4 ± 104.6 abc |
| Total glucans | ($\mu\text{eq g}^{-1} \text{ dw}$) | 1322.4 ± 235.6 abc | 1125.0 ± 144.6 bc |
| Hexose surplus | ($\mu\text{eq g}^{-1} \text{ dw}$) | 45.8 | |
|  | (%) | 20.0 |  |
| Agronomic surplus | ($\mu\text{eq g}^{-1} \text{ dw}$) | -123.1 | |
|  | (%) | -31.0 |  |
| CAM Particular 2b Condition (i.e. WZS2 in LC1) |  |  |  |
| Fractions | Units | H1 (Mean ± SE) | H2 (Mean ± SE) |
| Malic Acid | ($\mu\text{mol g}^{-1} \text{ dw}$) | 134.7 ± 26.6 c | 637.1 ± 278.4 a |
| Total sugars | ($\mu\text{eq g}^{-1} \text{ dw}$) | 1184.5 ± 16.0 bc | 1012.2 ± 136.3 cd |
| Sucrose | ($\mu\text{eq g}^{-1} \text{ dw}$) | 80.1 ± 27.2 cd | 241.3 ± 106.0 a |
| Glucose | ($\mu\text{eq g}^{-1} \text{ dw}$) | 403.3 ± 17.3 ab | 205.7 ± 15.4 d |
| Fructose | ($\mu\text{eq g}^{-1} \text{ dw}$) | 701.1 ± 20.1 abc | 565.2 ± 22.0 c |
| Total glucans | ($\mu\text{eq g}^{-1} \text{ dw}$) | 1579.7 ± 189.1 a | 1300.5 ± 118.5 abc |
| Hexose surplus | ($\mu\text{eq g}^{-1} \text{ dw}$) | 200.3 | |
|  | (%) | 80.0 |  |
| Agronomic surplus | ($\mu\text{eq g}^{-1} \text{ dw}$) | 53.5 | |
|  | (%) | 13.0 |  |

Equal letters indicate statistically non-significant differences ( $P > 0.05$ ). Comparisons are only valid for the same metabolite between H1 and H2. Data represent the mean ± standard error of three replicates. The hexose surplus is calculated based on the stoichiometric requirement for the amount of malic acid synthesized. The agronomic surplus is calculated using the highest malic acid value recorded in the experiment as the optimal CAM activity reference.

In CAM-particular2a (site TZ), hexose depletion from H1 to H2 followed the classical trend, yet it was insufficient to explain the observed malic acid accumulation. Specifically, 458.3 μmol·g^−1^ dw of malic acid were synthesized (571.4 μmol in H2 minus 113.1 μmol remaining in H1), while the decrease in hexoses (77.5 μeq·g^−1^ dw) could only account for 155 μmol of malic acid. The remaining 303.3 μmol (equivalent to 151.7 μeq of hexose) likely originated from starch hydrolysis. Climatic conditions for this event are presented in Table 2, Line 3.

In CAM-particular2b (site WZ), 502.4 μmol·g^−1^ dw of malic acid were synthesized (637.1 – 134.7), but the hexose decrease (172.3 μeq·g^−1^ dw) could account for only 344.6 μmol. Thus, 157.8 μmol (equivalent to 78.9 μeq of hexose) are again attributable to glucan hydrolysis. Table 2, Line 4, shows the corresponding environmental data.

The data clearly delineate the joint temperature and solar radiation ranges required for optimal CAM productivity in pineapple. While both variables must align to sustain high diel acid accumulation, suboptimal solar radiation—more frequent than low temperatures under tropical conditions—tends to be the limiting factor. This relationship will be explored in the Discussion with emphasis on its agronomic relevance.

### Condition CAM-particular3: minimal malic acid variation and nocturnal hexose increase

Table 5 presents CAM-particular3, characterized by low malic acid levels that remain nearly constant across the CAM cycle. Without pre-H1 values, it is not possible to assess prior decarboxylation. However, malic acid increased slightly from 109.4 μmol at H1 to 131.7 μmol at H2, indicating a nocturnal accumulation of only 22.3 μmol.

**Table 5.** CAM-particular3 condition results. Malic acid remains at low levels with minimal variation between H1 and H2, while total hexoses increase at H2. (Data correspond to the sample from CZS3 in LC1 and represent 10% of all samples in the experiment).

| CAM Particular 3 Condition |  |  |  |
| --- | --- | --- | --- |
| Fractions | Units | H1 (Mean ± SE) | H2 (Mean ± SE) |
| Malic Acid | ($\mu\text{mol g}^{-1}\text{ dw}$) | 109.4 ± 18.7 cd | 131.7 ± 21.7 cd |
| Total sugars | ($\mu\text{eq g}^{-1}\text{ dw}$) | 489.5 ± 206.4 e | 771.0 ± 307.6 de |
| Sucrose | ($\mu\text{eq g}^{-1}\text{ dw}$) | 141.3 ± 71.3 c | 137.8 ± 24.0 c |
| Glucose | ($\mu\text{eq g}^{-1}\text{ dw}$) | 50.2 ± 23.9 e | 272.7 ± 83.3 cd |
| Fructose | ($\mu\text{eq g}^{-1}\text{ dw}$) | 297.9 ± 118.3 d | 360.5 ± 254.7 d |

Equal letters indicate statistically non-significant differences ( $P > 0.05$ ). Comparison is only valid for the same metabolite across H1 and H2. Each value represents the mean ± standard error of three replicates.

On the other hand, the hexose amount increased from H1 to H2 by 281.5 μeq·g^−1^ dw. Although glucan data were not available for this particular case, it is reasonable to infer that the source of hexoses was chloroplastic starch hydrolysis. Climatic conditions during this sampling event— shown in Table 2, reference line 5—were atypical for pineapple, with low solar radiation and low temperatures.

It is plausible that under these limiting conditions, the plant did not reach Phase III of CAM, and instead remained in Phase II throughout the daylight hours. Consequently, nocturnal metabolism may have relied primarily on glycolysis, as part of a broader metabolic strategy to sustain cellular energy demands. The coordinated synthesis and degradation of starch, along with sucrose export, ensures that heterotrophic tissues continuously receive carbohydrates for metabolism and growth, both during light and dark periods (Griffiths et al., 2016).

This event also recorded the lowest soluble sugar content observed at the CZ site. Based on the net difference in hexoses between H2 and H1, the estimated nocturnal glycolytic activity was approximately 475 μeq hexose·g^−1^ dw. Considering the low starch reserves typically found under cold conditions, sustained exposure to similar weather might risk complete reserve depletion.

In discussing the circadian control of CAM, Hartwell (2007) noted that prolonged light exposure leads to an equilibrium in malate concentrations between the vacuole and cytoplasm, effectively inhibiting further malate accumulation due to feedback on PEP carboxylase (PEPc). Similarly, under constant darkness, Wilkins (1992) reported that malate accumulates in the cytoplasm during the PEPc-active phase, also inhibiting PEPc activity due to the lack of vacuolar sequestration. Under optimal conditions, vacuolar transport reactivates PEPc by removing cytoplasmic malate. However, energy deficits— such as those potentially induced by sustained darkness—could impair this transport, further suppressing PEPc function.

This may explain the minimal malic acid variation observed between H1 and H2 under CAM-particular3. Further investigation of this metabolic behavior under CZ-like conditions would benefit from a high-resolution, multi-day sampling protocol targeting the same plant and tissue, which is feasible from a logistical and technical standpoint.

### Analysis of individual metabolite profiles

The previous section presented the results of key CAM-related events and the overall dynamics of total hexoses (sum of sucrose, glucose, and fructose). Here, we examine the specific contribution of each sugar to the hexose pool.

Among the soluble sugars analyzed, sucrose stands out as the primary focus of physiological and agronomic interpretation, given its central role in phloem loading and long-distance transport of photosynthates. Moreover, the sampling strategy—targeting two critical time points of the CAM diel cycle (H1 and H2)—provided a favorable framework for exploring diel variation in sucrose concentration, both quantitatively and interpretatively. The Results section will therefore introduce the broader agronomic relevance of sucrose, which will be further developed in the Discussion. In contrast, glucose and fructose— despite being key intermediates—exhibit broader metabolic fates and more complex regulatory controls, which cannot be meaningfully resolved through the present two-point diel sampling. Consequently, their presentation in this section remains descriptive and is not followed by further interpretive analysis.

### Sucrose

The hexose contribution derived from sucrose generally ranged from the minimum detectable level—corresponding to the analytical method’s Limit of Detection (LoD)—to slightly above 500 μeq of hexose·g^−1^ dw. As shown in Figure 3, these minimum and maximum values are presented for agronomic reference and are categorized by each fixed effect examined in the study.

**Figure 3.**
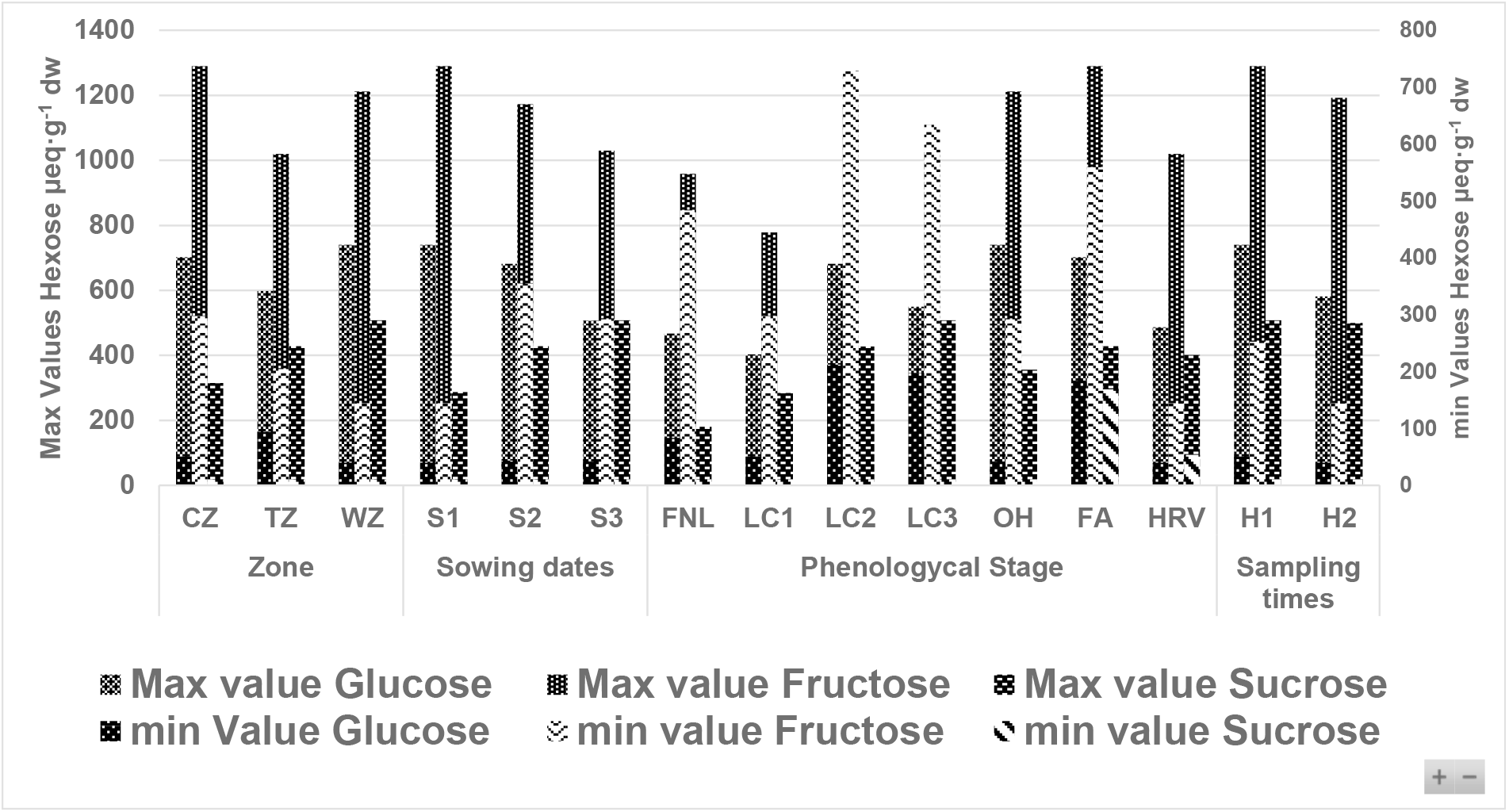
Minimum and maximum concentrations of glucose, fructose, and sucrose for each fixed effect: ecological zone, sowing date, phenological stage, and sampling time. Within each bar, values are shown in the following order: glucose, fructose, and sucrose. The left vertical axis corresponds to maximum values, while the right vertical axis corresponds to minimum values.

During the reproductive stages FA and HRV, sucrose levels followed a pattern of H1 > H2 at the WZ and TZ sites (see TZ-S2-FA, TZ-S2-HRV, WZ-S2-FA, and WZ-S2-HRV in Table 6). In contrast, at the CZ site, an opposite trend of H2 > H1 was observed, suggesting a distinct photosynthate acquisition and distribution pattern under colder climate conditions. Supporting environmental data—specifically solar radiation and temperature during these events—are presented in Table 2, line 7. The increase in sucrose levels at H2 for the CZ site is also evident in Table 6 (CZ-S2-FA and CZ-S2-HRV).

**Table 6.**
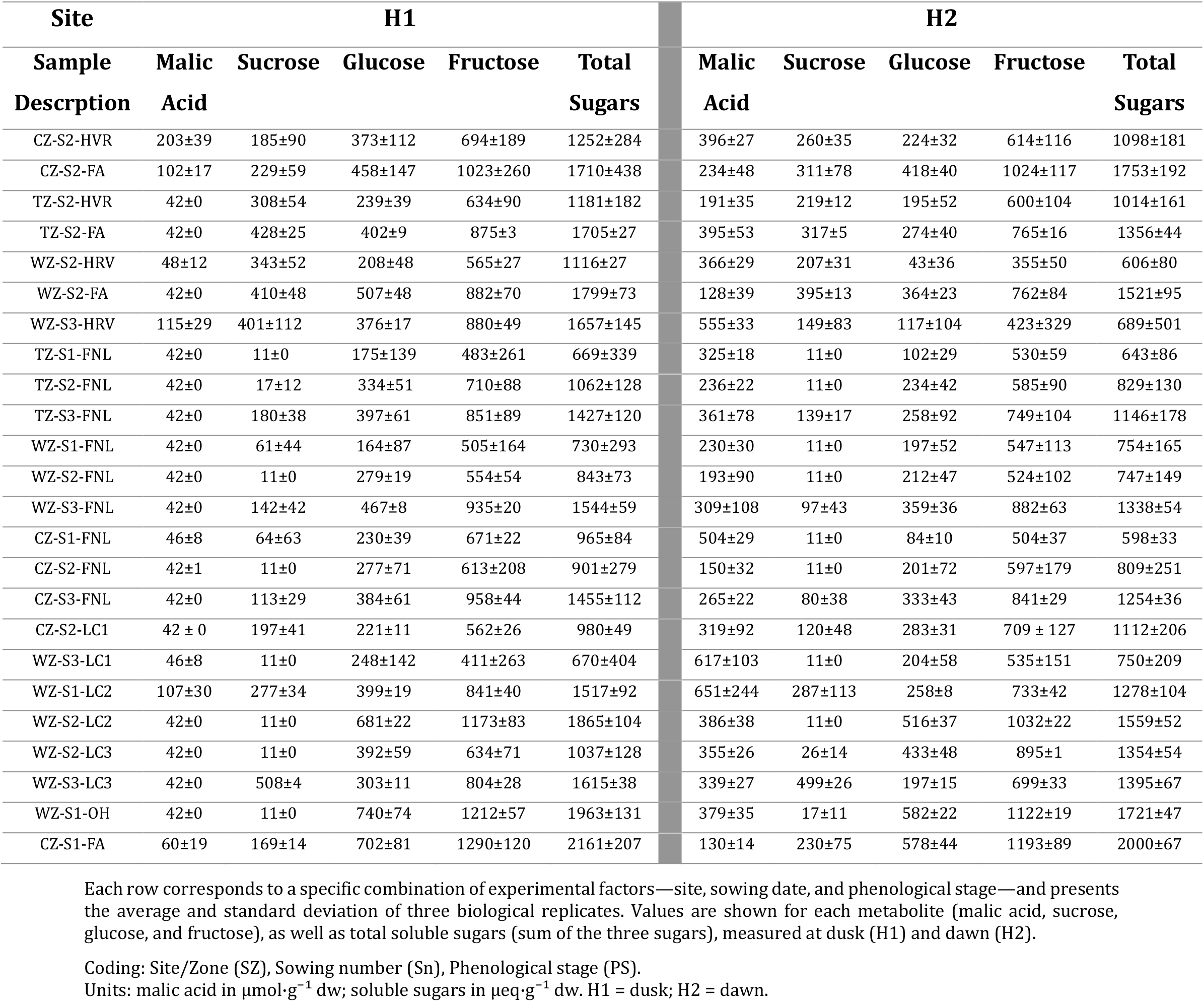
Representative examples of hexose and malic acid turnover in MD-2 pineapple leaves under contrasting experimental conditions.

The differences in sucrose dynamics between H1 and H2 reflect a modulation of carbon allocation priorities that appears contingent on both phenological stage (specifically some of the vegetative and reproductive phases previously described) and prevailing climate conditions. This pattern suggests a flexible regulatory mode, where export decisions shift between climate-driven control and phenology-driven demand, depending on the intensity and timing of developmental requirements and the occurrence of climatic constraints—not necessarily extreme—that limit carbon availability. These relationships will be further examined in the Discussion to elucidate their agronomic implications.

### Glucose

Glucose levels ranged from 40 to 740 μeq·g^−1^ dw across all samples (Figure 3). In TZ, glucose consistently declined from dusk (H1) to dawn (H2), consistent with the classical CAM pattern described by Carnal & Black (1989) (see Table 6, TZ). In contrast, at the WZ site, glucose increased overnight in events such as WZ-S1-FNL and WZ-S2-LC3, under high solar radiation and temperature (Table 2, lines 10 and 2), while sucrose remained undetectable or low at both time points. This suggests nocturnal glycolysis supporting sucrose synthesis and export, or alternatively, glucose accumulation via starch hydrolysis to meet metabolic demands during the night.

At the CZ site, only one event (CZ-S3-LC1) showed higher glucose at dawn (Table 5), with negligible changes in sucrose and malic acid levels under low radiation and temperature (Table 2, line 5). Although the glucose increase was observed at dawn, the low radiation and temperature during the preceding day likely restricted photosynthetic sugar production during Phases II–IV of CAM. As a result, carbohydrate availability during Phase I was constrained, limiting malic acid synthesis.

The absence of a dawn–dusk difference in sucrose levels at CZ may be explained by either a balance between export and synthesis during the night, or a temporary halt in sucrose export to restore reserves. In both scenarios, starch hydrolysis likely contributed to glucose accumulation. These data were collected under uncontrolled field conditions, and thus the physiological interpretations presented remain hypothetical. Further investigation into sugar export dynamics and their role in sustaining CAM and basal metabolism is warranted.

### Fructose

Fructose levels ranged from 144 to 1290 μeq·g^−1^ dw across all samples (Figure 3). Among the soluble sugars quantified, fructose consistently exhibited the highest concentrations at both dawn and dusk across all environments and phenological stages (see Table 6). This diel predominance was especially robust regardless of sampling location or leaf developmental phase.

This pattern has not been previously reported with comparable resolution under natural field conditions. Earlier studies in CAM plants described fructose accumulation during the day or its presence in specific tissues under controlled environments (Carnal & Black, 1989; Christopher & Holtum, 1996; Christopher & Holtum, 1998; Medina et al., 1993; Popp et al., 2003), but none provided field-based evidence of fructose being the most abundant soluble sugar throughout the diel cycle. Therefore, our data provide novel empirical support for this trait in pineapple operating under natural CAM.

Although the physiological consistency of high fructose levels is notable, its agronomic or diagnostic relevance remains unclear. These elevated concentrations may reflect fructose’s role as a primary metabolic intermediate in both sugar synthesis and starch mobilization.

### Malic acid

Malic acid content was consistently higher at dawn (H2) than at dusk (H1), clearly reflecting the constitutive CAM activity of pineapple plants (Tables 2, 4, 5, 6, and 8), as reported previously by Bartholomew (2018, 1982). Furthermore, the carbon isotope composition of pineapple flesh samples from all three study sites ranged from – 12.0‰ to –16.8‰. Because fruit development spans more than five months, these δ^13^C values provide an integrated record of carbon assimilation throughout the photosynthetic lifetime of the plant.

Carbon isotope composition has long been used to distinguish between plants with C3 versus C4 or CAM photosynthetic pathways (Bartholomew & Kadzimin, 1977). Since the enzyme Rubisco discriminates against ^13^C, C3 plants exhibit δ^13^C values typically ranging from –22‰ to –40‰, while C4 and CAM plants show less discrimination, yielding values between –9‰ and –19‰ (Subbarao & Johansen, 2001). In pineapple, δ^13^C values reported for leaves fall between –11‰ and –19‰ (Bartholomew & Malézieux, 1994), consistent with CAM metabolism and aligned with the values observed in our study. While these isotopic data do not distinguish CAM from C4 metabolism, they clearly separate C3 from CAM/C4 plants—a distinction relevant to this study.

Malic acid concentrations in the sampled leaves ranged from the method’s detection limit (41.6 μmol·g^−1^ dw, consistently at dusk) to nearly 800 μmol·g^−1^ dw at dawn. Within the framework of our standardized sampling protocol, the highest value recorded was 796 μmol·g^−1^ dw. To interpret this figure accurately, it is important to consider both the identity of the sampled leaf and the specific point on the leaf blade (see Methodology). Although not part of the formal dataset, we have independently recorded malic acid values exceeding 1300 μmol·g^−1^ dw in young leaves located at the crown or near the apex. These external observations are mentioned solely to clarify that our proposed reference of 796 μmol·g^−1^ dw as a potentially optimal agronomic CAM intensity corresponds strictly to leaves sampled following the standardized methodology, and should not be misinterpreted as a physiological upper limit for the species. Greater CAM activity in younger leaves has also been documented by Lin et al. (1994).

Despite the central role of malic acid in CAM, the diel patterns observed here reveal a complex and often inconsistent behavior that resists straightforward interpretation. While the measurements are robust, their agronomic relevance remains elusive when considered in isolation. Overall, our data underscore that relying solely on malic acid as a proxy for CAM function— particularly in field-grown pineapple—constitutes a limited and potentially misleading strategy. This issue will be revisited in the Discussion, where we argue for a broader, multi-metabolite approach to understand CAM plasticity in agronomic contexts

## 4. Discussion

Building upon the biochemical and isotopic data presented in the Results, the Discussion is organized thematically to emphasize physiological and agronomic insights rather than strictly following the methodological sequence. This approach reflects the exploratory nature of the study and prioritizes interpretative depth over a strict correspondence with the sequence of results. By focusing on findings with substantive interpretative value, the structure facilitates clarity, coherence, and relevance to applied pineapple cultivation.

### 1. CAM and agronomic reality

Although CAM has been widely characterized under controlled or semi-controlled conditions, its full expression under field conditions—and its consequences for agronomic performance— remain largely underexplored. In pineapple production, several common management practices show inconsistent or unpredictable outcomes that may reflect underlying physiological heterogeneity driven by CAM activity. The following three cases illustrate how such inconsistencies could be addressed through improved understanding of diel CAM dynamics in an agronomic context.

### Inconsistent degreening response

The application of ethephon to induce cosmetic changes in pineapple fruit displays substantial variability in efficacy, even under standardized technical conditions. This inconsistency has led to the adoption of multiple, often contradictory empirical practices (e.g., nighttime versus full-sun applications, among others), all reportedly successful in some cases, yet none consistently reproducible. A plausible explanation is that treatment efficacy depends on a dynamic internal metabolic state that is not externally observable— potentially linked to the diel phase of CAM metabolism. If malate accumulation, decarboxylation activity, and soluble sugar dynamics are not properly synchronized with the timing of application, the fruit’s responsiveness to ethephon may be diminished or erratic. External climate conditions may also contribute to this variability.

These empirical practices, widely circulated within the industry despite their inconsistent outcomes, underscore the urgent need for a mechanistic understanding that enables reliable standardization of degreening treatments.

### Burndown variability

A second critical practice in crop management is the application of herbicides to promote rapid biomass elimination and enable field renovation. This phase, known as burndown, is carried out when preparing for a new planting cycle and its effectiveness directly influences operational efficiency. In many cases, under technically equivalent conditions, the burndown response is highly variable, affecting the timing of equipment access to the field. Although the direct causes of this variability are not fully understood, the hypothesis that CAM metabolism may be involved gains relevance. Metabolic differences between plants—particularly in malate accumulation or sugar partitioning—could affect tissue response to herbicides and help explain why identically treated plots yield inconsistent results.

### Burned crowns during cold storage

Fruits with high malic acid content at the time of entering cold storage, due to incomplete remobilization of organic acids accumulated overnight, may be more susceptible to tissue damage. This phenomenon, known as “burned crowns,” typically affects only a fraction of exported fruit and occurs more frequently during the dry season. One likely explanation is that export containers often group fruits from different lots and harvest times, which masks the underlying physiological determinant. Understanding diel CAM dynamics in fruit tissue could inform agronomic adjustments— particularly in synchronizing harvest and precooling—to reduce the incidence of this type of damage.

These three cases reflect recurring field-level patterns suggesting that variability in agronomic performance may be associated with underlying CAM activity. However, their interpretation requires a more comprehensive understanding of diel CAM dynamics under **field conditions**— extending beyond isolated, single-day measurements—to pinpoint when and how metabolic transitions influence physiological responses. Additional cases, which will be addressed later in the discussion, further demonstrate how targeted physiological insights may inform critical management decisions in pineapple cultivation. These will be examined in the relevant sections based on their stronger association with specific diel CAM patterns or individual metabolites.

### 2. Four patterns of diel metabolism: evidence for CAM plasticity

The notion of a single, canonical CAM cycle in pineapple—based on diel organic acid dynamics under controlled conditions—has long shaped interpretations of its physiology. However, field-based evidence remains limited, especially under commercial tropical production environments where environmental fluctuations, long photoperiods, and sink activity significantly modulate carbon fluxes. In this study, we identified four distinct diel metabolic configurations in MD-2 plants grown under agronomically relevant conditions. The first of these was designated **CAM-classical**, due to its strong correspondence with traditional descriptions found in the controlled-condition literature.

The configurations designated CAM-particular 1 through 3 represent distinct expressions of diel CAM plasticity, identified during the analysis of field data collected under contrasting natural environments. These profiles were delineated based on recurring physiological patterns— differences in CAM intensity, carboxylation, decarboxylation, and mobilization of carbohydrate reserves—with some profiles observed across more than one environment. These findings challenge the adequacy of the classical CAM model for interpreting pineapple physiology under natural field conditions and underscore the need to reassess how CAM plasticity influences key agronomic traits such as source–sink balance, responsiveness to chemical treatments, floral induction, and harvest index.

### CAM-classical: reference profile consistent with canonical CAM

The CAM-classical profile, predominantly observed in plants from the WZ and TZ, closely mirrors the diel dynamics described in classical CAM literature (Bartholomew, 2018; Osmond, 1978), with significant nocturnal malate accumulation, daytime decarboxylation, and an increase in leaf sugar levels late in the day. This configuration was consistently associated with a range of maximum daytime temperatures of 30.1– 30.5 °C and a daily solar radiation of 11.1–19.8 MJ m^−2^ day^−1^, conditions commonly found in both sites (Table 1 and 2).

Physiologically, the CAM-classical profile allows for efficient carbon assimilation and daily sugar export, which is linked to the continuous vegetative growth observed in these plants. Agronomically, this is expressed in the production of wide, long leaves and greater overall plant biomass. However, plants exhibiting this profile tended to allocate assimilates primarily to expanding leaf tissue rather than accumulating stem reserves—an effect more pronounced in WZ, where leaf expansion was most vigorous.

This source–sink allocation pattern has direct implications for floral induction. Despite their high photosynthetic output, WZ plants exhibiting the CAM-classical profile showed resistance to natural flowering. This phenomenon suggests a physiological state in which carbon gain is decoupled from the plant’s readiness to transition to reproductive growth. In production zones such as WZ and in TZ during certain times of the year, where this pattern dominates, managing induction quality becomes critical, particularly in terms of synchrony and the number of fruitlets per stem. Consequently, plants with high vegetative biomass may produce a low number of fruitlets during forcing (Bartholomew & Sanewski, 2018), even though the plant possesses sufficient photosynthetic machinery to fill larger fruits. This highlights the importance of assessing the plant’s pre-forcing physiological condition.

### CAM-particular 1: A Metabolic Profile Associated with Resilience Under Environmental Stress

The CAM-particular 1 profile, observed in WZ plants, is distinguished by a metabolic dynamic where the complete decarboxylation of malic acid at dawn coexists with a notable increase in total hexose content at the end of the night (H2). This condition, which occurred in 9% of the cases, was recorded under environmental conditions of high solar radiation (between 15.6 and 19.7 MJ·m^−2^·day^−1^) and high maximum temperatures (32.3 to 33.4 °C). Leaf temperature data, with averages of 35-40°C and peaks of up to 75.7°C, are consistent with conditions that induce severe thermal stress (see WZ in Table 7).

**Table 7.** Air temperature, solar irradiance, and rainfall at the experimental sites during the metabolite measurements in MD-2 pineapple.

| Zone | Temperature $\delta$ | | | IrradianceΨ<br>Mj·m<sup>-2</sup>·day<sup>-1</sup> | Rainfallξ<br>mm |
| --- | --- | --- | --- | --- | --- |
|  | minimum | maximum | average |  |  |
|  | ----- °C ----- |  |  |  |  |
| CZ | 14.5 [9.3 – 17.0]**a** | 20.7 [16.8 – 24.2]**a** | 17.5 [14.6 – 20.3]**a** | 12.2 [4540 – 4378]**a** | 243 [2593 – 3256] |
| WZ | 21.3 [14.2 – 25.9]**b** | 33.1 [26.6 – 38.1]**b** | 26.9 [23.5 – 30.1]**b** | 19.5 [7305 – 6937]**b** | 137 [1394 – 1887] |
| TZ | 21.7 [15.1 – 24.3]**c** | 30.2 [22.4 – 34.4]**c** | 25.7 [20.7 – 28.9]**c** | 14.1 [5045 – 5263]**c** | 242 [2950 – 2859] |

$\delta$ Average of daily data over 2 years and, in square brackets, range minimum and maximum. $\psi$ Average of daily total over 2 years and, in square brackets, cumulative total (Year 1 and Year 2). $\xi$ Average monthly over 2 years and, in square brackets, cumulative total of Year 1 and Year 2. Equal letters within column represent non-significant differences for statistical test $p < 0.05$ .

The accumulation of hexoses in this profile is a manifestation of a high nocturnal rate of chloroplast starch hydrolysis. This massive carbohydrate mobilization, observed in genetically uniform plants with measurements normalized by dry weight, represents a plastic response that coincides with the greater foliar growth in WZ. The high availability of carbon during the night is a fundamental requirement for multiple anabolic processes, and this metabolic pattern is consistent with the larger leaf area and greater leaf dry weight observed in WZ compared to TZ and CZ (Table 8).

**Table 8.** Complementary biometric variables with significant allometric differences between agronomically relevant sites (TZ and WZ) during LC3.

|  | TZ |  |  |  | WZ |  |  |  |
| --- | --- | --- | --- | --- | --- | --- | --- | --- |
| Variable | S1 | S2 | S3 |  | S1 | S2 | S3 | IC 95% (±) |
| Leaf Area m² | 1.38 c | 1.69 b | 1.33 c |  | 1.80 ab | 1.84 ab | 1.93 a | 0.08 |
| Leaves dry weight g | 335.6 c | 367.0 bc | 261.7 d |  | 419.5 a | 377.8 b | 417.5 a | 17.2 |
| Stem dry matter % | 24 a | 20 b | 18 c |  | 20 b | 19 cb | 17 c | 0.01 |

Note: Statistical comparisons within each row. Different letters indicate significant differences ( $P < 0.05$ ); identical letters indicate no statistical difference.

This study demonstrates the coexistence of intensive carbohydrate mobilization with robust foliar physiology under conditions of extreme stress. To understand the underlying mechanism and validate the carbon allocation pathways, future studies are necessary to quantify lipid peroxidation and carbon flux towards the synthesis of proteins and lipids during the night.

### CAM-particular 2: an intermediate profile with a focus on hexose dynamics

The CAM-particular 2 profile, predominant in plants from the typical zone (TZ), presents a less pronounced nocturnal malate accumulation pattern compared to the classical profile (CAM-classical). Additionally, it is characterized by the presence of residual malate at the end of Phase IV (H1 sampling). This configuration was consistently observed on days with a range of maximum daytime temperatures of 28.5–31.2 °C and suboptimal daily solar radiation (<15 MJ m^−2^ day^−1^), below the threshold associated with high yields (Bartholomew et al., 2018; Malezieux et al., 1994). Alternatively, the residual malic acid at H1 might represent early nocturnal synthesis due to a premature onset of Phase I.

Physiologically, this profile suggests a reduced capacity for nocturnal carbon fixation compared to CAM-classical. However, the efficient sugar dynamics—characterized by a clear depletion of hexoses during the night—indicate that the plant maintains a high capacity to mobilize and export assimilates. The concurrent increase in sucrose from H1 to H2 in both cases suggests active sucrose synthesis for subsequent export. The stoichiometric and enzymatic implications of this pattern are examined in detail in Section 3. This type of climate is not unusual in TZ, especially during the rainy season.

If the low solar radiation conditions characteristic of the CAM-particular 2 profile persist for several days, carbohydrate reserves may become depleted, and plants would likely need to rely on alternative metabolic pathways to regenerate PEP. However, since these cloudy periods in TZ are interspersed with sunny days, these high radiation events are crucial for replenishing chloroplastic starch. This mechanism could be fundamental for maintaining high pineapple productivity in the humid tropics of Costa Rica and differentiates it from the consistently warm and high-radiation conditions of the dry tropics (WZ).

This preferential accumulation of dry matter in the stem is consistent with observations suggesting an explanation for the seasonality of natural flowering events in this zone.

### CAM-particular 3: a disrupted profile under low-temperature, low solar radiation conditions

The CAM-particular 3 profile was found predominantly in the Cold Zone (CZ), an experimental environment characterized by low temperatures and low solar radiation (Table 7). Physiologically, this profile is defined by an atypically low nocturnal malate accumulation, coupled with minimal variation in malate levels and an increase in hexose content at the end of Phase I (H2). This metabolic dynamic deviates markedly from the canonical pattern, and its low malate levels are consistent with a disruption in the CAM cycle.

The metabolic response observed in the CAM-particular 3 profile appears to be strongly associated with the environmental conditions of the CZ. The low solar radiation and temperature conditions observed in this zone are consistent with a restricted production of photosynthetic sugars during the day (Phases II–IV). This suggests that the availability of carbohydrates during the night was limited, which in turn may have restricted malate synthesis. This pattern of metabolic imbalance serves as an important contrast for understanding pineapple behavior in the Typical Zone (TZ). Our results suggest that if the low solar radiation conditions of the CAM-particular 2 profile were to be sustained, the CAM metabolism in the TZ could converge toward an altered expression similar to that of the CAM-particular 3 profile. Therefore, the analysis of this profile is crucial for defining the limits of CAM plasticity in pineapple and for highlighting the importance of the climatic conditions in the TZ, where the intermittence of sunny and cloudy days prevents this type of metabolic disruption.

From an agronomic perspective, the findings in the CZ provide essential insights into the effects of suboptimal climatic conditions. In this environment, plants exhibited slower growth, although they showed stem dry matter accumulation greater than 30% (compared to <20% in TZ and WZ). This suggests that while CAM can confer resilience under stress conditions, the suboptimal expression of the metabolic cycle under certain environmental conditions can be linked to undesirable agronomic outcomes, such as a disruption of vegetative growth and a predisposition to natural flowering.

### 3. Agronomic interpretation of hexose balance and CAM efficiency: a comparison with Carnal & Black (1989)

From an agronomic perspective, the research by Carnal & Black (1989) made a significant contribution by reporting the total balance of hexose turnover—including soluble sugars and glucans—rather than limiting the analysis to CAM cycle sufficiency. Although their primary objective was biochemical, their dataset inadvertently provides information of agronomic interest: a quantitative surplus of assimilated hexoses. Such a balance is seldom reported in CAM research, which typically focuses on the biochemical sufficiency for completing the diel acid cycle, rather than evaluating the availability of assimilates for structural growth or export. According to their analysis, there was an estimated surplus of approximately 40%. A small portion of this surplus was attributed to respiration, though the precise fraction was not quantified. The remainder was inferred to support biosynthesis, yet this component remains the least documented in the literature (Walker et al., 2021).

Using the same logic as Carnal & Black (1989), we calculated the hexose surplus and surplus (%) values based on our data from Tables 1, 3, and 4. Additionally, we estimated agronomic surplus and agronomic surplus (%) using a benchmark for high CAM performance. Specifically, we propose an exploratory reference of 796 μmol of malic acid·g^−1^ dw—the highest value recorded in our field data—as a potential agronomic target for CAM intensity. Carnal & Black reported photosynthetic metabolite concentrations on a fresh weight basis, which we converted to dry weight using the average moisture content (84.09 ± 0.06%) obtained from 717 samples in our dataset. Under this conversion, the surplus of hexoses reported by Carnal & Black (115 μeq·g^−1^ dw) corresponds to 44% of their observed malic acid accumulation. However, when benchmarked against our proposed agronomic target, this surplus becomes negative (–20.5 μeq·g^−1^ dw; – 5%).

The agronomic surplus and percentage surplus values from Carnal & Black (1989) can be compared to our data in Table 3, though they originate from different baselines of malic acid production and hexose turnover. Their nocturnal malate production (525 μmol·g^−1^ dw) closely matches our data in Table 4 (CAM-particular2b, 502.4 μmol·g^−1^ dw), while their reduction in soluble sugars from dawn to dusk (261.3 μeq·g^−1^ dw) is comparable to our Table 1 result (237.7 μeq·g^−1^ dw).

Comparing the environmental conditions of the CAM-classical event (Table 1, TZ site) with CAM-particular1 (Table 3, WZ site) reveals that the former aligns more closely with climate conditions typically associated with high pineapple productivity (Bartholomew et al., 2018; Hepton et al., 1993). The greatest hexose availability for export and malic acid synthesis was observed at the TZ site. This supports the idea that the CAM-classical pattern is more agronomically favorable.

Climatic conditions in TZ and WZ (in Table 4) were thermally optimal for high productivity but showed suboptimal solar radiation (<15 MJ·m^−2^·day^−1^), below the threshold linked to high yields (Bartholomew et al., 2018; Bartholomew & Malézieux, 1994). These data suggest that both radiation and temperature must align to achieve maximal CAM performance.

Our dataset confirms that these are typical CAM conditions. Nonetheless, Tables 1, 3, and 4 show considerable variability in CAM productivity. The best indicator of high photosynthetic performance is not solely the amount of malate produced at night or sugars accumulated during the day, but the concentration of glucans—primarily starch— stored in chloroplasts. In fact, the most substantial physiological divergence between our field data and the results of Carnal & Black lies in glucan content. Our findings indicate that chloroplastic glucans function as the principal carbon reservoir for subsequent sucrose synthesis and export via starch hydrolysis, positioning them as a key determinant in the agronomic efficiency of CAM in pineapple.

According to Carnal & Black (1989), the replenishment of soluble sugar reserves during the day—primarily glucose and fructose— supports malic acid accumulation at night through glycolysis and the activity of PPi-dependent phosphofructokinase (PPi-PFK), which catalyzes the conversion of fructose-6-phosphate to fructose-1,6-bisphosphate and promotes PEP formation. Their study emphasized the energetic advantage of this pathway in the CAM cycle but did not directly investigate the role of ATP-dependent PFK (ATP-PFK). Moreover, environmental variables such as temperature or solar radiation were not reported, making it unclear whether their observations apply broadly across different agroecological contexts.

Conversely, during periods of persistently low solar radiation (Table 5, Table 2, line 5), glucose and fructose levels increased from dusk to dawn, while sucrose declined slightly—even though malic acid accumulation remained limited. These patterns suggest that hexoses were generated via hydrolysis of chloroplastic glucans but were not efficiently converted into malate. According to Carnal & Black (1989), this inefficiency could reflect a shift in glycolytic flux. The enzyme ATP-dependent phosphofructokinase (ATP-PFK), which relies on full glycolysis to generate PEP, operates more slowly than the alternative enzyme that uses pyrophosphate (PPi-PFK), which can utilize soluble hexoses directly. We hypothesize that under favorable CAM conditions in pineapple, PPi-PFK serves as the primary enzyme for nocturnal PEP production. In contrast, when environmental conditions inhibit CAM intensity, ATP-PFK may become the predominant pathway—less efficient and unable to sustain high malate synthesis.

### 4. Malate is not enough: the limitations of single-marker approaches

In previous sections, we documented that malic acid was not always fully decarboxylated at dusk (e.g., Table 4). Several physiological and environmental factors may account for this incomplete decarboxylation:

Leaf area and self-shading: During early developmental stages such as FNL and LC1, reduced leaf area allows for more complete decarboxylation. As the canopy expands, however, self-shading increasingly limits photosynthesis in intermediate and lower leaves, with the extent of this limitation depending on solar radiation levels, often resulting in residual malic acid at the end of Phase IV.

Temperature and solar radiation: The highest concentration of residual malic acid at dusk was recorded at Site CZ, particularly under conditions of low solar radiation and temperature (Table 5; see also climate data in Table 2, line 5). In these conditions, malate was not fully decarboxylated by the end of Phase IV. Residual malic acid also persisted under high temperature and solar radiation, which may induce photoinhibition. For example, in WZ-S1 during LC2, residual malic acid was still detectable at the end of Phase IV (Tables 2 and 6). Photoinhibition in CAM plants has been reported primarily during Phase IV, although Lüttge (2001) noted it may also occur in Phase III. However, this possibility lacks experimental confirmation and appears physiologically unlikely, especially in pineapple, where high internal CO_2_ concentrations during Phase III are expected to mitigate photoinhibition. A complementary study on thermal time calibration conducted as part of this research showed improved developmental prediction when temperature thresholds above 33 °C were excluded from the model—supporting the interpretation that CAM performance declines under excessive heat and solar radiation (Vásquez-Jiménez et al., 2024).

Microclimatic heterogeneity within the canopy: Variations in light exposure caused by overlapping or adjacent leaves can generate localized differences in CAM activity within a single leaf. In pineapple, this heterogeneity is a structural feature of the plant: leaves are arranged in a full 360° around the stem, and each one experiences a distinct light regime throughout the day. Consequently, the variability in leaf photosynthetic production is a common factor for the entire plant. Popp et al. (2003) likewise reported differences in metabolite content across leaf sections, associated with chlorophyll gradients.

Shifting focus to carbohydrate dynamics, high malic acid accumulation at the end of CAM Phase I was commonly associated with reductions in glucose and fructose concentrations from dusk to dawn. In contrast, sucrose levels tended to remain stable or declined less markedly, suggesting preferential use of glucose and fructose in nocturnal PEP regeneration (e.g., Table 4, CAM-particular condition 2b). This pattern aligns with the mechanism proposed by Carnal & Black (1989), who demonstrated that under CAM conditions, sucrose synthesized in the cytosol is primarily directed toward phloem export rather than malate production.

### 5. Metabolic economy and functional roles of glycolytic pathways

The three agronomic scenarios presented previously—the inconsistent response to fruit degreening, the variability in burndown, and the incidence of cold-storage crown damage— illustrate how a lack of deep understanding of CAM metabolism in the field creates operational challenges and economic losses. These limitations cannot be resolved simply by measuring malic acid synthesis or titratable acidity, as commonly done in CAM studies under field conditions — an approach that overlooks diel complexity and metabolic flexibility. The deep-rooted reliance on malic acid concentration as the primary indicator of CAM activity has been a cornerstone of plant physiology, particularly in controlled studies (Luttge & Ball, 1987; Osmond, 1978), but its insufficiency to explain variability in pineapple under field conditions is now evident.

Our findings challenge the sufficiency of this approach by demonstrating that robust malate dynamics do not always correlate with an efficient and balanced use of carbon. The particular metabolic profiles we identified reveal a more complex landscape where malate fixation, sugar recycling, and carbon partitioning are not always perfectly synchronized.

For example, in the CAM-particular 1 profile, we observed a full decarboxylation of malic acid, a pattern typically interpreted as optimal CAM function. However, this was accompanied by a significant nocturnal surplus of hexoses, a finding that suggests a possible decoupling between malate synthesis and the metabolic mobilization of sugars. Similarly, in the CAM-particular 2 profile, hexose consumption was insufficient to account for the total malate produced. This result, which challenges the model of Carnal & Black (1989), implies that the plant relied on additional carbon sources—likely chloroplastic starch—to sustain nocturnal fixation.

These observations highlight that CAM efficiency is not solely a function of how much carbon is fixed at night, but of how the available carbohydrate pool is managed. Therefore, a more complete view must integrate the dynamics of hexoses, sucrose, and glucans to accurately assess the plant’s functional status. Based on this, malate may serve as an indicator of cycle activity, but carbohydrate dynamics are the true reflection of the plant’s metabolic economy. Depending exclusively on malate as a single diagnostic marker risks misinterpreting the plant’s physiological health, which could lead to suboptimal agronomic decisions and perpetuate the variability in fruit degreening, burndown, and cold damage.

By analyzing pineapple sugar metabolism, specifically through diel patterns of soluble sugar accumulation and turnover, our results provide evidence about the energy economy underlying CAM photosynthesis. Pineapple employs two alternative glycolytic pathways, each with distinct energetic costs, indicating that CAM metabolism modulates not only acid accumulation but also the energy expenditure required for carbon recycling.

These findings extend the classical characterization of CAM beyond its well-known roles in water use efficiency and acid storage, highlighting it as a system integrating both carbon and energy fluxes. Future work should aim to quantify these pathways’ relative contributions under variable field conditions to further elucidate their ecological and agronomic significance.

### 6. CAM and agronomic reality: insights from spatial metabolic variability in pineapple

#### 6a. Sucrose as a physiological indicator of seed quality

In young plants at the FNL stage, sucrose levels in the leaves of S1 and S2 plantings—characterized by seeds with high stem starch content—were generally low and approached the Limit of Detection (LoD) at both dusk and dawn (see Table 6 for FNL values across TZ, WZ, and CZ sites), suggesting limited sucrose availability at this stage. It has been documented that pineapple seeds with elevated stem starch levels support vigorous early growth (Vásquez-Jiménez et al., 2018). In contrast, the higher sucrose content observed in the leaves of S3 plants may reflect a different carbon allocation strategy: seeds with lower initial stem starch content may mobilize photosynthate from leaves to the stem via sucrose transport to compensate for limited stored reserves. Sucrose plays a key role in phloem loading and long-distance translocation (Bavnhøj et al., 2023; Liu et al., 2012; Stein & Granot, 2018; Tong et al., 2022), supporting the accumulation of starch in the stem for root initiation and early leaf expansion.

By the end of the FNL stage, leaves appear to replenish the chloroplastic starch reserves previously exported during establishment. However, interpreting stem dry matter concentration requires caution. Seeds that took longer (in both thermal and chronological time) to reach the target fresh weight often accumulated greater stem dry matter. While this favors rapid establishment, it also increases the risk of natural flowering (NF). Conversely, seeds with lower stem dry matter, but that reached the planting threshold more quickly, may be agronomically preferable depending on sowing timing and NF risk.

Although a standardized protocol was applied to maintain uniform seed characteristics in each planting, unavoidable variations in sowing dates introduced differences in physiological backgrounds. While seed type, weight, and size were controlled, the age of the source plants used to produce the vegetative seed differed across sowings—reflecting current industry practices in seed management and maintained as such in this study.

At the V-stage FNL, our data are consistent with the hypothesis that photosynthetic and glycolytic sucrose (quantified in leaves) could serve as metabolic indicators of sucker seed quality, warranting further validation. This opens the door to a research protocol aimed at validating whether easily distinguishable seed types—selected a priori using field-accessible criteria, such as external morphology and fresh weight, and, importantly, blocked by physiological age estimated via growing degree days—exhibit consistent and physiologically meaningful differences in carbon metabolism. While such a protocol would necessarily involve detailed sampling and metabolite analysis under controlled comparative trials, its ultimate goal is not to implement metabolite profiling as a routine tool in agronomic decision-making. Rather, it aims to empirically test whether seed selection strategies that combine current agro-industrial standards with physiological age stratification can reliably predict metabolic readiness and planting quality. In this framework, metabolic analysis serves strictly as a validation tool, not as an operational requirement. This strategy has the potential to refine current agro-industrial paradigms, which prioritize seed uniformity primarily by weight, by integrating a physiologically grounded layer of selection. A central aim is to improve fruit uniformity at harvest—an enduring weakness of the current seed selection paradigm, in which sowing uniformity rarely translates into uniform fruit development.

The higher sucrose levels observed in S3-FNL (Table 6: WZ-S3-FNL and TZ-S3-FNL) are likely related to the younger age of the seedplot from which the S3 sucker seeds were obtained. Although sucrose differences between sowings disappear after LC1, this early variation may still impact seed quality and uniformity in commercial pineapple production.

Higher leaf sucrose levels during FNL likely reflect an active export of photosynthate from the leaves to the stem, particularly in seeds with initially low starch reserves. In this context, sucrose functions as a primary transport molecule, mobilizing carbon to support root initiation and early development. This mechanism is consistent with the observed patterns in the S3 sowing, where lower stem reserves may have triggered early carbon allocation via sucrose.

#### 6b. Sucrose dynamics during reproductive stages and harvest index

Sucrose concentrations were frequently close to the LoD during early phenological stages, including at dawn (H2). However, this pattern changed markedly during the FA and HRV stages, where minimum sucrose values were consistently higher (Figure 3), and dusk-to-dawn fluctuations became much more pronounced—with turnover rates exceeding 60% in specific cases. Notable examples include WZ-S2-HRV and TZ-S2-FA (Table 6), with the most extreme case occurring in WZ-S3-HRV under high solar radiation and temperature (Table 2, line 6).

We propose that these dynamics reflect the plant’s developmental phase. In FA and HRV, the fruit enters its main filling phase and becomes the strongest sink in the plant. To sustain this demand, sucrose is actively exported from the leaves, intensifying source–sink carbon redistribution. This results in significant sucrose turnover between dusk and dawn, especially under favorable environmental conditions. This behavior aligns with established source–sink relationships in pineapple, as described by Bartholomew et al. (2018) and Smith et al. (2018), where the fruit attracts the majority of photoassimilates in late stages.

At the WZ and TZ sites, during LC1, sucrose levels at H2 were higher than at H1, although absolute concentrations remained low (Table 4), under solar radiation and temperature conditions below 12 MJ·m^−2^·day^−1^ and 30 °C, respectively (Table 2, lines 3–4). In contrast, at CZ—also during LC1— where temperatures were below 20 °C and solar radiation reached only 5 MJ·m^−2^·day^−1^, sucrose levels were high and remained stable between H1 and H2 (Table 2, line 8; Table 5), suggesting elevated sink demand.

The highest absolute sucrose concentrations were also associated with elevated solar radiation. For example, WZ-S3-LC3 reached 508.1 ± 3.9 μeq of hexose·g^−1^ dw during H1, following three consecutive days with solar radiation above 19 Mj·m^−2^·day^−1^ (Table 6; Table 2, line 9). Although the WZ site generally exhibited the highest sucrose values (Table 6), the TZ site displayed the greatest net hexose accumulation across the CAM cycle.

This suggests that while sucrose synthesis responds positively to high radiation and temperature, total carbon accumulation is also modulated by broader source–sink balance and site-specific metabolic adjustments.

Our data differ from the findings of Medina et al. (1993), who reported that only sucrose consistently exhibited an inverse relationship with the titratable acidity cycle. In our case, sucrose dynamics do not explain malic acid accumulation. While we cannot definitively exclude the participation of sucrose in glycolysis and the nocturnal formation of PEP, our results show either an increase or minimal variation in sucrose from H1 to H2, accompanied by substantial reductions in glucose and fructose—an observation also reported by Carnal & Black (1989). The decline in total hexose is largely attributed to these two monosaccharides, supporting the hypothesis that glucose and fructose, rather than sucrose, serve as the primary respiratory substrates for PEP formation during the night. Conversely, sucrose concentrations appear to follow a pattern more consistent with nocturnal export. This is physiologically coherent, as sucrose is a key transport form of carbon: being chemically stable and not redox-active, it can be mobilized through the phloem without prior biochemical transformation (Hernández-Bernal et al., 2022).

In earlier stages, high sucrose concentrations are not observed because there is no strong sink demanding large amounts of photoassimilates— except in FNL, as previously discussed. However, even in FNL, sucrose levels remain lower than those in reproductive stages due to the limited size of the photosynthetic apparatus and the plant’s low metabolic capacity. At this stage, the plant has only recently been established and is still undergoing acclimatization (Vásquez-Jiménez et al., 2023).

It has been documented that, following floral induction, dominant apical growth is interrupted, leading to a marked decrease in the demand for photosynthate for leaf expansion, while the developing inflorescence initially exerts only a minor sink influence (Bartholomew, 2018). As a result, dry matter—primarily starch— accumulates in the stem, since the leaves continue producing more photoassimilates than are required by the limited sinks at this stage.

This physiological condition is climate-dependent. In sites with favorable conditions for high productivity, such as TZ, this response pattern is expected. However, in our data from the OH reproductive stage, there was no consistent evidence of active sucrose export despite maximal leaf area and low sink demand. This suggests that sucrose allocation toward stem reserves may follow a regulated pathway, likely involving continuous export derived from chloroplastic glucan hydrolysis during both day and night (see Figure 1). Alternatively, early reproductive sinks such as the developing fruit may already begin receiving photoassimilates in the form of sucrose, explaining the increased leaf sucrose levels observed from the FA stage onward.

We hypothesize that in plants at the TZ and WZ sites during the reproductive stages FA and HRV, sucrose export occurred continuously—both during the day (from photosynthesis) and at night (from the hydrolysis of starch reserves in chloroplasts; see Figure 1). In contrast, at the CZ site, sucrose production appeared to depend more on starch and glucan hydrolysis, likely reflecting the lower suitability of this environment for CAM metabolism (see a general comparison of the environments in Table 7). This is supported by the increase in total soluble sugars from H1 to H2 at CZ (Table 5), indicating a predominance of nocturnal carbohydrate mobilization rather than daytime photosynthetic activity.

Given the limited agronomic literature on CAM expression and phloem export dynamics in pineapple, this interpretation should be considered tentative pending further experimental confirmation. However, it integrates consistent patterns observed across contrasting climates and phenological stages. For instance, during reproductive stages (R-Stages), sucrose concentrations tend to be higher at H1 than at H2 in TZ and WZ, whereas the opposite pattern is observed in CZ—suggesting differential regulation of sugar export under less favorable CAM conditions.

From the FA stage onward, the H1 vs. H2 pattern appears to depend on whether the climate supports effective CAM function. In environments favorable to CAM (like WZ and TZ), the intense production of photosynthetic sucrose during the day results in higher H1 than H2 values—even if some starch is hydrolyzed at night. Conversely, in climates where CAM activity is restricted (such as CZ), sucrose tends to be higher in H2 due to reliance on stored carbohydrate mobilization (Borland et al., 2014; Hartwell, 2007).

In WZ during vegetative stages, sucrose may fluctuate above or below glucose depending on local climatic conditions. At TZ, however, glucose consistently exceeded sucrose across all vegetative samples. A similar pattern was observed in CZ, with the exception of CZ-S3-LC1 in H1, where sucrose briefly surpassed glucose. This was corrected at H2, with glucose again exceeding sucrose levels (Table 5), reinforcing the pattern of nocturnal sugar remobilization supported by CAM-related carbohydrate dynamics (Borland et al., 2014; Carnal & Black, 1989).

Considering the negative impact of natural flowering on pineapple production, it is noteworthy that no natural flowering was observed at the WZ site during our study period. This raises the question of whether a higher proportion of sucrose relative to glucose at key moments—such as the end of each phyllotaxic leaf cycle—could indicate a reduced risk of floral induction.

In vegetative stages (V-Stages), it was consistently observed that when sucrose levels at dusk (H1) were close to or equal to the limit of detection (LoD), they remained equally low at dawn (H2), while glucose and fructose levels were consistently high (e.g., WZ-S3-LC1, WZ-S2-LC2, WZ-S1-OH in Table 6). This pattern may indicate limited or reduced sucrose synthesis activity under these specific developmental or environmental conditions. While this condition was still observed in the R-Stage OH, it became absent from the FA stage onward. As previously discussed, the allocation of photosynthate toward fruit filling in reproductive stages requires significant resources. High temperatures and solar radiation stimulate lush foliage in pineapple, producing numerous, broad, and flaccid leaves with shallow grooves (Bartholomew, 2018). This morphology implies a high demand for sucrose, which is most efficiently supplied through the direct export of photosynthetic hexoses. Since MD-2 does not produce new vegetative structures other than leaves during V-Stages (Vásquez-Jiménez et al., 2023), the elevated sucrose contents observed in V-Stage samples at the WZ site could help explain the greater leaf area recorded in this zone (see Table 8).

Recently, questions about the transport and partitioning of photosynthates in long-established crops—such as grapevine—have regained relevance, particularly regarding their agronomic implications (Walker et al., 2021). These authors emphasized that addressing such questions requires examining the anatomical structure and compartmentalization of immediate photosynthate storage within the leaf. The fact that this basic physiological aspect remains a topic of research in crops cultivated for millennia highlights the agronomic value of analyzing these dynamics in tropical species like pineapple. In this context, foliar sugar profiles, particularly sucrose, may offer key insights into carbon mobilization and allocation, with potential applications for physiological understanding and agronomic decision-making.

The elevated glucose levels in TZ contribute to sucrose production, but the demand for and transport of carbohydrates to storage organs occurs at a slower, more sustained rate, rather than being driven by rapid structural growth. At the WZ site, sucrose production is typically high during daylight under intense solar radiation and temperature, whereas in TZ, sucrose production is more discrete but continuous throughout the day and night. This pattern may explain the higher stem dry matter content observed at TZ (see Stem dry matter in Table 8), as well as the higher harvest index recorded in this site—consistently above 0.75—compared to WZ, where it averaged below 0.70. This is consistent with the observations of Hepton et al. (1993), who noted that in warmer tropical environments, dry matter is preferentially allocated to vegetative growth, particularly to leaf development, thereby reducing the harvest index. In contrast, under cooler conditions, assimilates tend to be temporarily stored as non-structural dry matter in the stem, which can later be remobilized to support fruit development. As a result, plants with lower vegetative mass may produce relatively heavier fruits, thus increasing the harvest index.

Significant starch accumulation occurs in the leaf and stem of pineapple plants in subtropical areas, but very little or no starch (less than 10% dry matter) is found in tropical environments (Bartholomew, 2018). This observation also aligns with the occurrence of NF. Naturally induced plants have a smaller leaf area and higher dry matter content in the stem compared to neighboring plants that are not induced at the time of NF. For instance, plants that were not naturally induced at the TZ site in V-Stage LC3 resulted in less than 20% stem dry matter at the time of forcing. When plants showed signs of NF between LC2 and LC3, the LC2 sampling revealed that they had up to 30% dry matter at the stem level and a leaf area of only 0.9 m^2^. Increases in stem dry matter due to the start of flowering have been reported by Bartholomew (2018).

Our data suggest that the export of sucrose synthesized daily from soluble photosynthetic hexoses in sites like WZ is predominantly allocated to the development of broad and long leaves. As a result, there is limited transport of sucrose toward the stem for dry matter accumulation. This reflects a physiological trade-off in phloem allocation: only one of the two sink pathways—structural expansion or reserve accumulation—appears to dominate. Consequently, when sucrose is directed primarily to leaf growth, natural flowering (NF) does not occur, consistent with observations showing increased stem dry matter prior to NF.

When artificially inducing flowering in WZ-like environments, it is critical to ensure high induction quality—defined as the number of fruitlets per stem (Bartholomew & Sanewski, 2018). If only a few fruitlets are initiated, the harvest index decreases, resulting in small and commercially undesirable fruits, despite the plant’s large vegetative size and robust photosynthetic capacity.

Although the photosynthetic apparatus in sites like WZ can supply sufficient assimilates for fruit growth, these agroecosystems often exhibit resistance to artificial induction, resulting in poor fruitlet formation. While both induction factor and induction quality have been conceptually defined—as previously noted—the industry typically focuses only on the former, neglecting quality, which more directly affects fruit size.

## 5. Conclusions

The results of this study demonstrate that the Crassulacean Acid Metabolism (CAM) syndrome in pineapple is far more metabolically and agronomically complex than classical CAM models suggest. This complexity arises from the plasticity of CAM expression, which is modulated by environmental conditions, spatial variability, and the plant’s developmental stage, factors that have been largely overlooked in previous studies focused on limited biochemical markers such as malic acid concentration alone.

Our findings show that reliance on malate as the sole indicator of CAM activity risks oversimplification and may lead to misinterpretation of physiological status, especially under field conditions where carbohydrate dynamics—particularly hexose and sucrose metabolism—play essential roles in carbon allocation and energy management. The identification of distinct diel metabolic profiles reveals that carbon partitioning between malate fixation, sugar recycling, and starch reserves varies significantly depending on environmental context and metabolic demand, reflecting a finely tuned balance between carbon conservation and energy expenditure.

In warm tropical environments, sucrose accumulation at the end of the CAM cycle appears as a key feature supporting rapid vegetative growth through phloem transport, while in cooler or less favorable climates, plants increasingly rely on stored glucan reserves to sustain nocturnal carbon fixation. Fructose consistently emerges as the most abundantly synthesized soluble sugar, indicating its central role in the nocturnal respiratory metabolism linked to PEP generation and malate synthesis.

Moreover, spatial variability in carbohydrate metabolism among sites and sowing types highlights the agronomic importance of seed physiological age and source plant characteristics in determining metabolic readiness and subsequent growth performance. These insights underscore the need for seed selection strategies that incorporate physiological markers beyond traditional morphological or weight-based criteria, aiming to enhance uniformity and yield in commercial pineapple production.

The study also expands the conceptual understanding of CAM by elucidating the functional significance of alternative glycolytic pathways—namely, ATP-dependent and pyrophosphate-dependent phosphofructokinases—which modulate energy consumption in carbohydrate recycling and malate synthesis. This metabolic flexibility allows pineapple to optimize its energy economy in response to fluctuating environmental stresses, further emphasizing the complexity of CAM as a dynamic system integrating carbon and energy fluxes.

Advancing agricultural management and breeding of CAM-dependent crops such as pineapple requires a holistic approach that integrates diel metabolic profiling, environmental variability, and physiological seed quality. Such integration will enable more precise agronomic interventions, improved crop uniformity, and resilience under diverse growing conditions. Future research should prioritize quantifying the relative contributions of key metabolic pathways in field conditions to deepen our understanding of CAM plasticity and its ecological and agronomic implications.

## 6. Acknowledgements

We thank Dr. Giovanni Saenz for the initial coordination with researchers from Universidad Nacional de Costa Rica and MICITT that made this research possible.

We thank Dr. Thomas Guzman for providing one experimental plot and laboratory equipment at the Instituto Tecnológico de Costa Rica to perform part of this research.

We thank the company BASF of Costa Rica, especially Agr. Eng. Alvaro Chinchilla, for facilitating the experimental plot and various logistical tasks essential for this research.

Finally, we thank Agr. Eng. Oscar Mario Vargas for facilitating the experimental plot of the cold zone, which was essential for conducting this research.

## 7. Funding sources

This work was partially supported by MICITT (Ministry of Science, Innovation, Technology and Telecommunications of Costa Rica) [grant number MICITT-PINN-CON-620-2019].

## 8. Conflict of interest

The authors declare that they have no conflict of interest.

## Notes

### Competing Interest Statement

The authors have declared no competing interest.

